# Discovery of Hierarchical Representations for Efficient Planning

**DOI:** 10.1101/499418

**Authors:** Momchil S. Tomov, Samyukta Yagati, Agni Kumar, Wanqian Yang, Samuel J. Gershman

## Abstract

We propose that humans spontaneously organize environments into clusters of states that support hierarchical planning, enabling them to tackle challenging problems by breaking them down into sub-problems at various levels of abstraction. People constantly rely on such hierarchical presentations to accomplish tasks big and small – from planning one’s day, to organizing a wedding, to getting a PhD – often succeeding on the very first attempt. We formalize a Bayesian model of hierarchy discovery that explains how humans discover such useful abstractions. Building on principles developed in structure learning and robotics, the model predicts that hierarchy discovery should be sensitive to the topological structure, reward distribution, and distribution of tasks in the environment. In five simulations, we show that the model accounts for previously reported effects of environment structure on planning behavior, such as detection of bottleneck states and transitions. We then test the novel predictions of the model in eight behavioral experiments, demonstrating how the distribution of tasks and rewards can influence planning behavior via the discovered hierarchy, sometimes facilitating and sometimes hindering performance. We find evidence that the hierarchy discovery process unfolds incrementally across trials. We also find that people use uncertainty to guide their learning in a way that is informative for hierarchy discovery. Finally, we propose how hierarchy discovery and hierarchical planning might be implemented in the brain. Together, these findings present an important advance in our understanding of how the brain might use Bayesian inference to discover and exploit the hidden hierarchical structure of the environment.

## Introduction

Imagine you have a sudden irresistible craving for your favorite ice cream that is only made by a boutique ice cream shop in Lugo, Spain. You must get there as soon as physically possible. What would you do? When faced with this unusual puzzle, most people’s first response is that they will look up a flight to Spain. When asked what they would do next, most people say that they would order a taxi to the airport, and when questioned further, that they would walk to the taxi pickup location. Importantly, nobody says or even thinks anything like “I will get up, turn left, walk five steps, etc.”, or even worse, “I will contract my left quadricep, then my right one, etc.”. This example illustrates hierarchical planning (Figure 1): people intuitively reason at the appropriate level of abstraction, first sketching out a plan in terms of transitions between high-level *states* (in this case, countries), which is subsequently refined in progressively lower levels and specific steps (Wiener and Mallot, 2003). This is often done in an online fashion, with details being resolved on-the-fly as the plan is being executed (for example, you would not ponder what snack to buy at the airport before actually getting there). This ability of humans (Balaguer et al., 2016) and animals (Geddes et al., 2018) to organize their behavior hierarchically allows them to flexibly pursue distant goals in complex environments, even for novel tasks that they may have never encountered previously.

**Figure 1.**
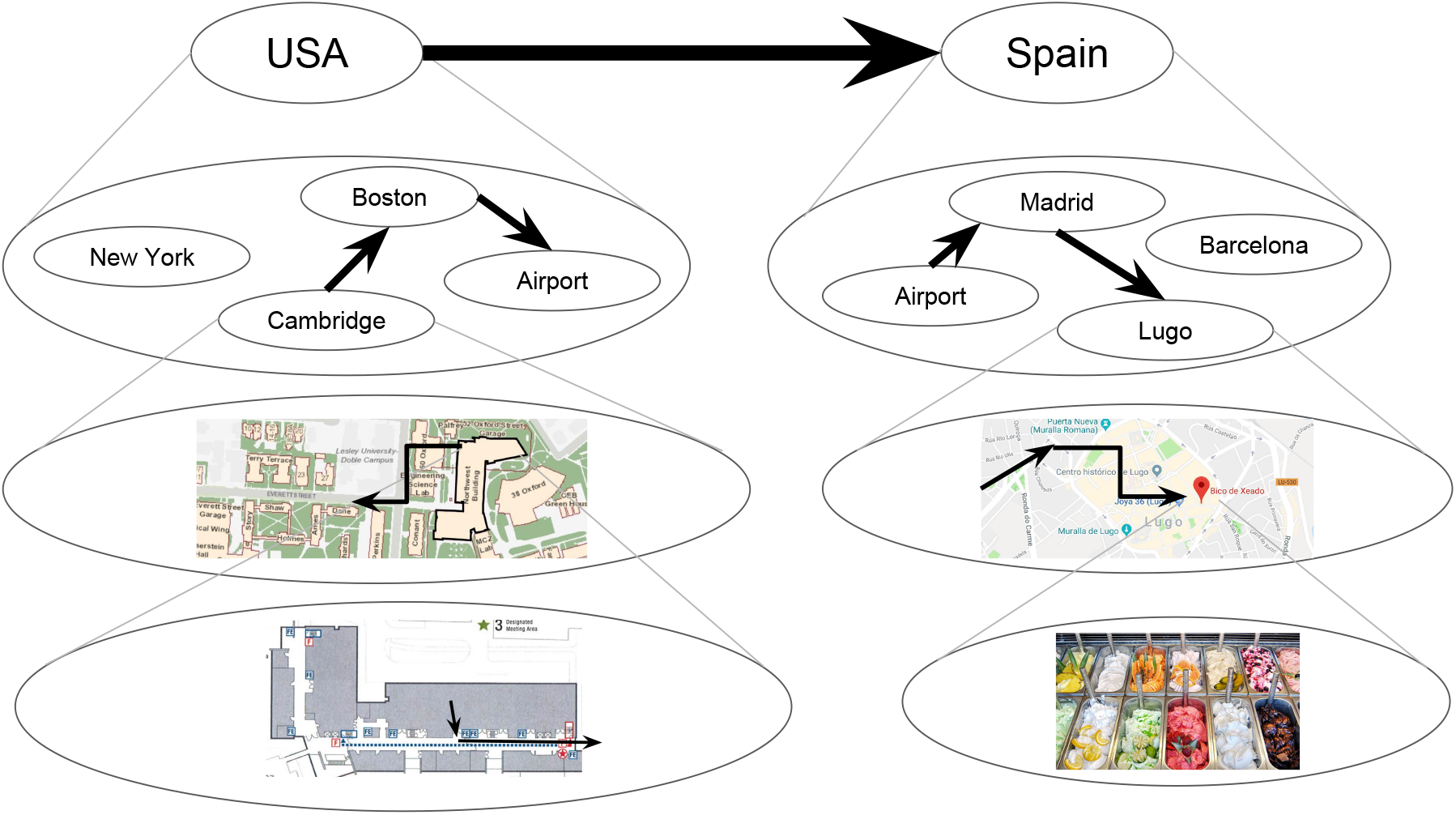
Example of hierarchical planning. How someone might plan to get from their office in Cambridge to their favorite ice cream shop in Lugo, Spain. Circles represent states and arrows represent actions that transition between states. Each state represents a cluster of states in the lower level. Thicker arrows indicate transitions between higher-level states, which often come to mind first.

The study of hierarchical behaviors has deep roots in psychology and neuroscience (Lashley, 1951). Much work has been done to characterize the emergence of such behaviors in humans and animals, often focusing on the acquisition of temporally extended action sequences or *action chunks* that unfold over different time scales. Action chunking occurs after extensive training, involves a specific set of brain regions, and is thought to be essential for pursuing long-term goals and planning (Graybiel, 1998; Smith and Graybiel, 2013).

Yet in order to leverage action chunks for planning, an agent must also be equipped with a hierarchical representation of the environment, with clusters of states or *state chunks* representing parts of the world at different levels of abstraction. In our ice cream example, the current country, city, neighborhood, building, room, and location within the room are all valid representations of the agent’s current state and are all necessary in order to plan effectively. Correspondingly, research shows that people spontaneously discover hierarchical structure (Wiener and Mallot, 2003; Schapiro et al., 2013), that the discovered hierarchies are consistent with formal definitions of optimality (Solway et al., 2014a), and that people use these hierarchies to plan (Balaguer et al., 2016). Some of these studies have uncovered distinct neural correlates of *high-level states*, which are sometimes referred to as state clusters, communities, contexts, abstract states, or state chunks (Schapiro et al., 2013; Balaguer et al., 2016). Research has also begun to uncover the neural correlates of hierarchical planning and action selection (Balaguer et al., 2016; Ribas-Fernandes et al., 2011). Yet despite these advances, the computational mechanisms underlying the discovery of such hierarchical representations remain poorly understood.

In this study, we propose a Bayesian model of hierarchy discovery for planning. Drawing on the structure learning literature (Gershman et al., 2015) and on concepts developed in robotics (Fernández and González, 2013), the model provides a normative account of how agents with limited cognitive resources should chunk their environment into clusters of states that support efficient planning. The central novel contributions of this paper are both empirical and theoretical. Our main empirical contribution is to show that the distribution of tasks (experiments one through five) and rewards (experiments six and seven) in the environment can influence the inferred state chunks, whereas past studies have focused exclusively on the effects of environment topology. Our main theoretical contribution is to unify these and previous findings under a single normative model that explains why these phenomena occur, and that encompasses a class of process models that could be leveraged to investigate the implementational details of state chunking in the brain.

In simulations one through five, we demonstrate that the model accounts for previously reported behavioral effects of the environment topology, such as detection of transitions across state clusters (Schapiro et al., 2013), identification of topological bottlenecks (Solway et al., 2014a), preference for routes with fewer state clusters (Solway et al., 2014a), and slower reaction times to transitions that violate topological structure (Lynn et al., 2018). Additionally, the model makes specific predictions about the way in which the distributions of tasks and rewards constrain hierarchy discovery, which in turn constrains planning. We test these novel predictions in a series of eight behavioral experiments. Experiment one shows that the distribution of tasks encountered in the environment can induce different state clusters, even when the topological structure of the environment does not promote any particular clustering. Experiment two shows how this in turn could lead to either improved or hampered performance on novel tasks. Experiment three examines the progression of inferred hierarchies as a function of which tasks participants have seen so far and reveals that participants are sensitive to changes in both the uncertainty and the mode of the posterior distribution over hierarchies. Experiments four and five replicate the results of experiments one and two in a fully visible environment, showing that the effects cannot be explained by incorrect inferences about topological structure. Experiment six shows how rewards generalize within state clusters, while experiment seven shows how rewards can induce clusters that constrain planning even in the absence of state clusters. Finally, experiment eight demonstrates that people explore in a way that maximally reduces uncertainty about the hierarchy, implying that people consider a probability distribution over hierarchies rather than a single hierarchy. Together, these results provide strong support for a Bayesian account of hierarchy discovery, according to which the brain implicitly assumes that the environment has a hidden hierarchical structure, which is rationally inferred from observations.

## Theoretical Background

The question of how agents should make rewarding decisions is the subject of reinforcement learning (RL; Sutton and Barto, 2018). With a long history crisscrossing the fields of psychology, neuroscience, and artificial intelligence, RL has made major contributions to explaining many human and animal behaviors (Rescorla et al., 1972), the neural circuits underlying these behaviors (Schultz et al., 1997), and allowing artificial agents to achieve human-level performance on tasks that were previously beyond the capabilities of computers (Mnih et al., 2015). Approaches rooted in RL therefore offer promising avenues to understanding decision making and planning.

Most RL algorithms assume that that an agent represents its environment as a Markov decision process (MDP), which consists of states, actions and rewards that obey a particular conditional independence structure. At any point in time, the agent occupies a given state, which could denote, for example, a physical location such as a place in room, a physiological state such as being hungry, and abstract state such as whether a subgoal has been achieved, or a complex state such as a conjunction of such states. The agent can transition between states by taking actions such as moving or eating. Some states deliver rewards, and it is assumed that the agent aims to maximize reward. In an MDP, the transitions and rewards are assumed to depend only on the current state and action.

Following previous authors (Solway et al., 2014a; Balaguer et al., 2016), we assume for simplicity that states are discrete and that transitions between states are bidirectional and deterministic (although these restrictions could be relaxed; see General Discussion). In this case, states and actions could be represented by an undirected graph *G* in which the vertices correspond to states and the edges correspond to actions. We will use the graph *G* in place of the transition function *T* that is traditionally used to characterize MDPs.

In graph theoretic notation, *G* = (*V, E*), where

- *V* is the set of vertices (or nodes) such that each node *v* ∈ *V* corresponds to a state that the agent can occupy, and
- *E*: {*V* × *V*} is a set of edges such that each edge (*u, v*) ∈ *E* corresponds to an action that the agent can take to transition from state *u* to state *v* or vice versa.

In the following analysis, we use the terms node and state interchangeably. We also treat edges, actions, and transitions as equivalent. For simplicity, we restrict our analysis to unweighted graphs, which is equivalent to assuming that all actions carry the same cost and/or take the same amount of time (our analysis extends straightforwardly to weighted graphs; see General Discussion). We also assume the agent has learned G, which is equivalent to model-based RL in which the agent learns the transition function.

The task of planning to get from a starting state *s* to a goal state *g* efficiently can thus be framed as finding the shortest path between nodes *s* and *g* in *G*. This is a well-studied problem in graph theory and the optimal solution in this setup is the breadth first search (BFS) algorithm (Cormen et al., 2009). BFS works by first exploring the neighbors of s, then exploring the neighbors of those neighbors, and so on until reaching the goal state g. States whose neighbors haven’t been explored yet are maintained in an active queue, with states getting removed from one end of the queue as soon as their neighbors are added to the other end. Intuitively, this corresponds to a forward sweep that begins at s and spreads in all directions across the edges until reaching g, akin to the way in which water might spread in a network of pipes. Its simplicity and performance guarantees have made BFS a standard tool for planning in classical artificial intelligence (Russell and Norvig, 2016).

However, the time and memory that BFS requires is proportional to the number of states (assuming an action space of constant size) since the size of the active queue and the potential length of the plan grow linearly with the size of the state space. Formally, the time and memory complexity of BFS is *O*(*N*), where *N* = |*V*| is the total number of states. In environments in which real-world agents operate, this number can be huge; in the realm of navigation alone, there could be billions of locations where a person has been or may want to go. Assuming that online computations such as planning involve systems for short-term storage and symbol manipulation, this far exceeds the working memory capacity of people who can only accommodate a few items (Miller, 1956; note that we assume the graph is already stored in a different system for long-term storage of relational information, such as the hippocampus). Furthermore, even without such working memory limitations, artificial agents such as robots would still take a long time to plan the route before they can take the first step. When using a naive or “flat” representation in which the agent plans over low-level actions (for example, individual steps or even joint actuator commands), computing a plan is at least as complicated as actually executing it (Figure 2A), and in reality the complexity could be much larger.

**Figure 2.**
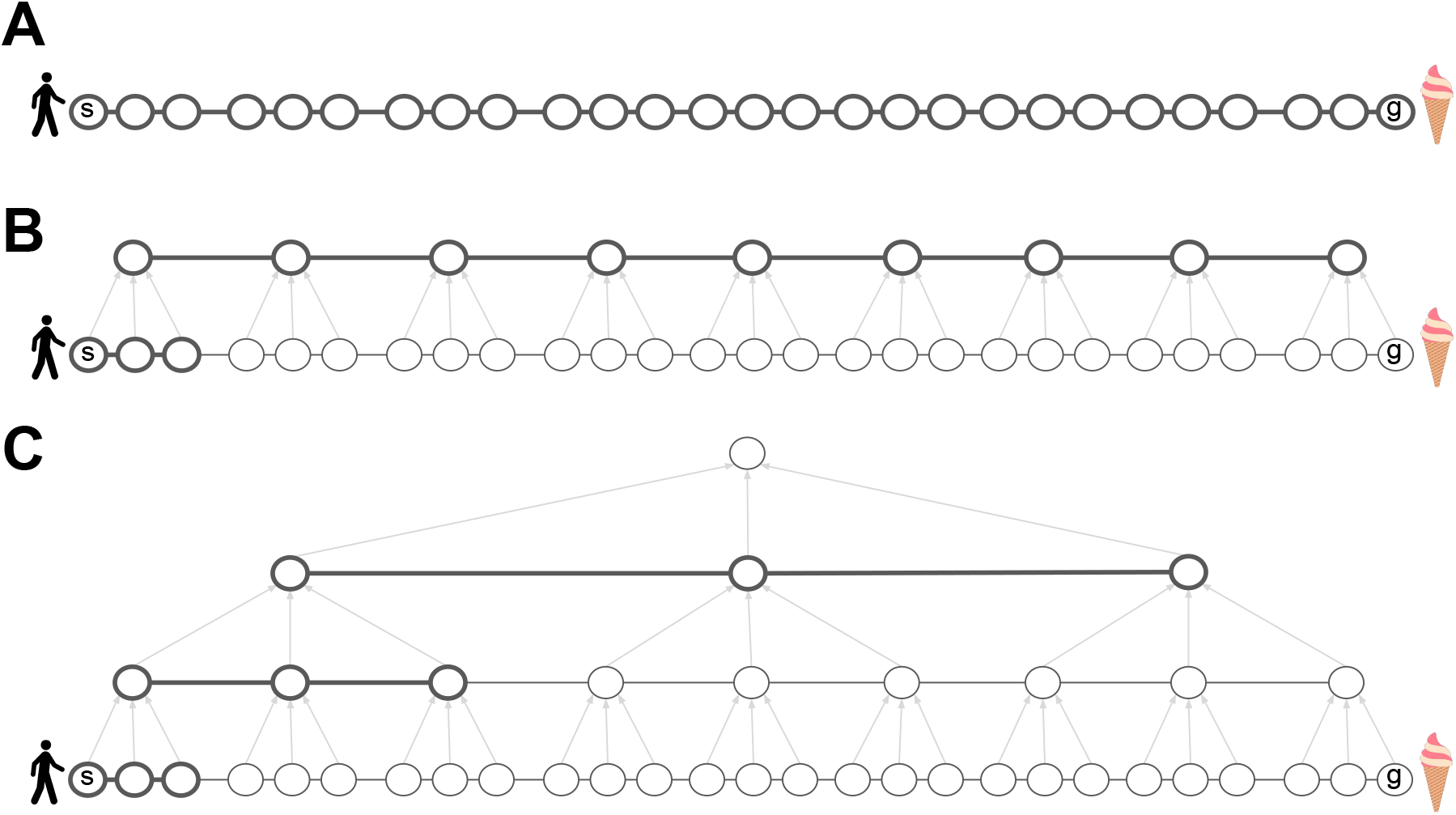
Hierarchical representations reduce the computational costs of planning. A. Planning in the low-level graph *G* takes at least as many steps as actually executing the plan. All nodes and edges are thick, indicating that they must all be considered and maintained in short-term memory in order to compute the plan. B. Introducing a high-level graph *H* alleviates this problem. At any given time during plan execution, the agent only needs to consider the high-level path and the low-level path leading to the next cluster, recomputing the latter on-the-fly. Gray arrows indicate cluster membership. C. The hierarchy can be extended recursively, further reducing the time and memory requirements of planning.

To overcome this limitation, work in the field of robotics has led to the development of data structures and algorithms for hierarchical planning (Fernández and González, 2013). Similar ideas have been put forward in other fields; see General Discussion. The key idea is that an agent can group neighboring states from the flat low-level graph *G* into state clusters (state chunks), with each cluster represented by a single node in another graph *H* (the high-level graph), which will be smaller and hence easier to plan in. When tasked to get from state *s* to state *g* in *G*, the agent can first plan in the high-level graph *H* and then translate this high-level plan into a low-level plan in *G*. Crucially, after finding the high-level path in *H*, the agent only needs to plan in the current state cluster in *G*, that is, it only needs to plan how to get to the next cluster (Figure 2B), and then repeat the process in the next cluster, and so on, until reaching the goal state in the final cluster.

This can drastically reduce the working memory requirements of planning, since the agent only needs to keep track of the (much shorter) path in *H* and the path in the current state cluster in *G*. Importantly, this also reduces planning time, allowing the agent to begin making progress towards the goal without computing the full path in *G* – the agent can now follow the high-level plan in *H* and gradually refine it in *G* on-the-fly, during execution. This approach can be applied recursively to deeper hierarchies in which high-level states are clustered in turn into even higher-level states, and so on until reaching a single node at the top of the hierarchy that represents the entire environment (Figure 2C). Planning using such a hierarchical representation can be orders of magnitude more efficient than “flat” planning, and also accords with our intuitions about how people plan in everyday life.

A particular instantiation of this form of hierarchical planning is the hierarchical breadth first search (HBFS) algorithm, which is a natural extension of BFS (see Appendix). It can be shown that with an efficient hierarchy of depth *L* (that is, consisting of *L* graphs, with each graph representing clusters in the lower graph), the time and memory complexity of HBFS is 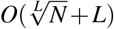 (Fernández andGonzález, 2013). Thus in a graph with *N* = 1,000,000,000 states and a hierarchy *L* = 9 levels deep, HBFS will only require on the order of 19 memory and time units to compute a plan, compared to 1,000,000,000 time and memory units for BFS, or any other flat planner. Note that while executing the plan would still take *O*(*N*) time, HBFS quickly computes the first few actions in the right direction, and can then be applied repeatedly to keep computing the following actions in an online fashion. While it may seem that this hierarchical scheme simply transfers the burden to the long-term storage system, which now needs to remember *L* graphs instead of one, it can be shown that the storage requirements of an efficient hierarchical representation are *O*(*N*) (Fernández and González, 2013), comparable to those of a flat representation of G alone. Following previous authors, (Botvinick et al., 2009; Solway et al., 2014a), we restrict our analysis to *L* = 2 levels for simplicity, noting that our approach extends straightforwardly to deeper hierarchies (see General Discussion).

However, in order to reap the benefits of efficient planning, the hierarchical representation necessarily imposes a form of lossy compression. In particular, each successive graph in the hierarchy loses some of the detail present in the lower level graph. This could lead to hierarchical plans that correspond to suboptimal paths in *G*. For example, in Figure 3A, there is a direct edge that can take the agent from starting state *s* to go state *g* in a single action. However, since the edge is not represented by the high-level graph *H*, HBFS will compute a detour through the state cluster *w*, akin to real-life situations in which people prefer going through a central location rather than taking an unfamiliar shortcut.

**Figure 3.**
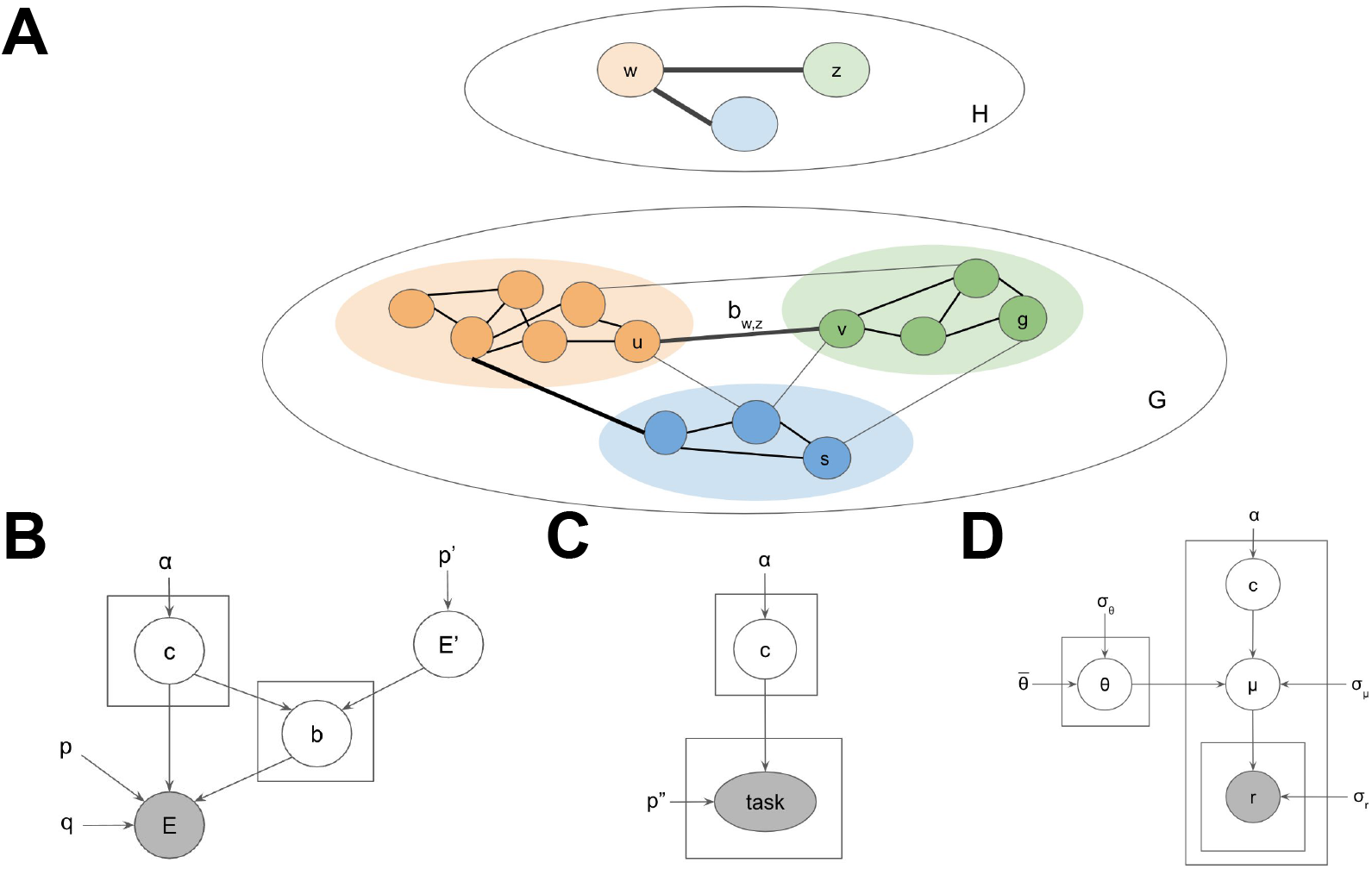
Generative model for environments with hierarchical structure. A. Example low-level graph *G* and high-level graph *H*. Colors denote cluster assignments. Black edges are considered during planning. Gray edges are ignored by the planner. Thick edges correspond to transitions across clusters. The transition between clusters *w* and *z* is accomplished via the bridge *b_w,z_* = (*u, v*). B. Generative model defining a probability distribution over hierarchies *H* and environments *G*. Circles denote random variables. Rectangles denote repeated draws of a random variable. Arrows denote conditional dependence. Gray variables are directly observed by the agent. Uncircled variables are constant. C. Incorporating tasks into the generative model. The rest of the generative model is omitted for clarity. D. Incorporating rewards into the generative model. The rest of the generative model is omitted for clarity.

Since finding the shortest path will not always be possible, some hierarchies will tend to yield better paths than others. The challenge for the agent is then to learn an efficient hierarchical representation of the environment that facilitates fast planning across many tasks, without placing an overwhelming burden on working memory and long-term memory systems. How agents accomplish this in the real world is the central question of this paper, to which we propose a solution in the following section.

## A Bayesian model of hierarchy discovery

Our proposal assumes two main computational components:

1. An online planner that can flexibly generate plans and select actions on-demand with minimal time and memory requirements.
2. An off-line (and possibly computationally intensive) hierarchy discovery process that, through experience, incrementally builds a representation of the environment that the planner can use.

The focus of this paper is the second component, which must satisfy the constraints imposed by the first component. For the first component, we use HBFS in order to link the hierarchy to behavior, noting that any generic hierarchical planner will make similar predictions. Note that we assume only the online planner is constrained by working memory limitations and time demands – as in the ice cream example, the high-level sketch of a plan is often computed within seconds of the query. In contrast, the hierarchical representation that supports this computation – a rich abstraction of the world, with knowledge of particular locations that belong to cities that belong to countries connected by flights – has been refined through years of experience and is deeply ingrained in long-term memory.

One approach to deriving an optimal hierarchy discovery algorithm would be to define the constraints of the agent, such as memory limitations and computational capacity, its utility function, and the constraints of the environment, such as the expected structure and tasks. Here we adopt an alternative approach, motivated by the literature on structure learning which has been used to successfully account for a wide range of phenomena in the animal learning literatire (Gershman et al., 2015). The key idea is that the environment is assumed to have a hidden hierarchical structure that is not directly observable, which in turn constrains the observations the agent can experience. The agent can then infer this hidden hierarchical structure based on its observations and use it to plan efficiently. Assuming that some hierarchies are *a priori* more likely than others, this corresponds to a generative model for environments with hierarchical structure, which the agent can invert to uncover the underlying hierarchy based on its experiences in the environment.

Formally, we represent the observable environment as an unweighted, undirected graph *G* = (*V, E*) and the hidden hierarchy as *H* = (*V′, E′, c, b, p′, p, q*), where:

- *V* is the set of low-level nodes or states, corresponding to directly observable states in the environment,
- *E*: {*V* × *V*} is the set of edges, corresponding to possible transitions between states via taking actions,
- *V′* is the set of high-level nodes or states, corresponding to clusters of low-level states,
- *E′*: {*V′* × *V′*} is the set of high-level edges, corresponding to transitions between high-level states,
- *c*: *V* → *V′* are the *cluster assignments* linking low-level states to high-level states,
- *b*: *E′* → *E* are the *bridges* which link high-level transitions back to low-level transitions,
- *p′* ∈ [0, 1] is the density of the high-level graph,
- *p* ∈ [0, 1] is the within-cluster density of *G*,
- *q* ∈ [0, 1] is the across-cluster density penalty of *G*.

Together, (*V′, E′*) define the high-level graph, which we also refer to as *H* for ease of notation. Each low-level state *u* is assigned to a cluster *w* = *c_u_*. Each high-level edge (*w, z*) has a corresponding low-level edge (the bridge) (*u, v*) = *b_w,z_*, such that *c_u_* = *w* and *c_v_* = *z* (see Figure 3A). Bridges (sometimes referred to as bottlenecks or boundaries) thus specify how different clusters are connected in *H*. Bridges are the only cross-cluster edges considered by the hierarchical planner; all other edges between nodes in different clusters are ignored, leading to lossy compression that improves planning complexity but could lead to suboptimal paths. In contrast, all edges within clusters are preserved. The purpose of the cluster assignments *c* is to translate the low-level planning problem in *G* into an easier high-level planning problem in *H*. The purpose of the bridges is to translate the solution found in *H* back into a low-level path in *G*. For a detailed description of how HBFS uses the hierarchy to plan, see Appendix. For simplicity, we only allow a single bridge for each pair of clusters, noting that our approach generalizes straightforwardly to multigraphs (i.e., allowing multiple edges between pairs of nodes in *E* and *E′*) and maintains its performance guarantees as long as the maximum degree of each node is constant (that is, the size of the action space is *O*(1)).

Informally, an algorithm that discovers useful hierarchies would satisfy the following desiderata:

1. Favor smaller clusters.
2. Favor a small number of clusters.
3. Favor dense connectivity within clusters.
4. Favor sparse connectivity between clusters (Cowan, 2001), with the exception of the bridges that connect clusters.

Intuitively, having too few (for example, one) or too many clusters (for example, each state is its own cluster) creates a degenerate hierarchy that reduces the planning problem to the flat scenario, and hence medium-sized clusters are best (desiderata 1 and 2). Additionally, the hierarchy ignores transitions across clusters, which could lead to suboptimal paths generated by the hierarchical planner. It is therefore best to minimize the number of cross-cluster transitions (desiderata 3 and 4). The exception is bridges, which correspond to the links between clusters.

These desiderata can be formalized as a generative model for hierarchies and environments (Figure 3B):

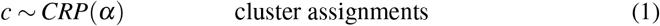

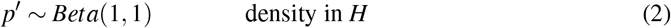

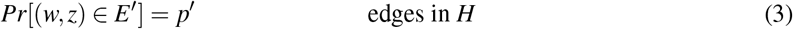

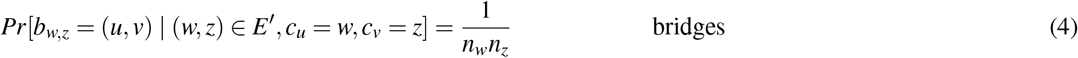

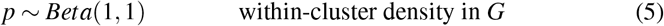

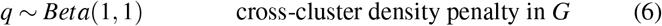

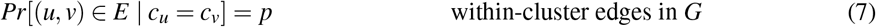

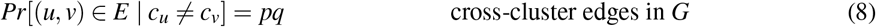

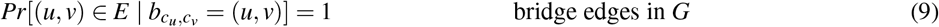

Where *n_w_* = |{*u*: *c_u_* = *w*}| is the size of cluster *w* and CRP is the Chinese restaurant process, a nonparametric prior for clusterings (Gershman and Blei, 2012).

Eq. 1 fulfills desiderata 1 and 2, with the concentration parameter *α* determining the trade-off between the two: lower values of *α* favor few, larger clusters, while higher values of *α* favor more, smaller clusters. Eq. 2 and Eq. 3 generate the high-level graph *H*, with higher values of *p′* making *H* more densely connected. Eq. 4 generates the bridges by connecting a random pair of nodes (*u, v*) for each pair of connected clusters (*w, z*). Eq. 5 and Eq. 7 fulfil desideratum 3 by generating the low-level edges in *G* within each cluster, with higher values of *p* resulting in dense within-cluster connectivity. Eq. 6 and Eq. 8 fulfill desideratum 4 by generating the low-level edges across clusters, with higher values of *q* resulting in more cross-cluster edges. Note that *pq* < *p* and hence the density of cross-cluster edges will always be lower than the density of within-cluster edges. Finally, Eq. 9 ensures that each bridge edge always exists.

This generative model captures the agent’s subjective belief about the generative process that gave rise to the environment and the observations. This belief could itself have been acquired from experience or be evolutionarily hardwired. The assumed generative process defines a joint probability distribution *P*(*G, H*) = *P*(*G*|*H*)*P*(*H*) over the observable graph *G* and the hidden hierarchical structure *H* that generated it. Importantly, the generative process is biased to favor graphs *G* with a particular “clustered” structure. In order to discover the underlying hierarchy *H* and to use it to plan efficiently, the agent must “invert” the generative model and infer *H* based on *G*.

Formally, hierarchy discovery can be framed as performing Bayesian inference using the posterior probability distribution over hierarchies *P*(*H*|*G*):

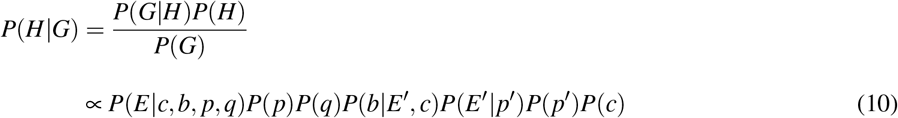

This predicts that states which are more densely connected will tend to be clustered together. We assess this prediction in simulations one through five.

### Task distribution

Previous studies have demonstrated that people discover hierarchies based on topological structure (simulations one through five). However, other factors may also play a role. In particular, the distribution of tasks that an agent faces in the environment may make some hierarchies less suitable than others (Fernández and González, 2013), independently of the graph topology. For example, if the agent has to frequently navigate from state *s* to state *g* in the graph *G* in Figure 3A, then the current hierarchy *H* would clearly be a poor choice, even if it captures the topological community structure of *G* well. Since hierarchical planning is always optimal within a cluster, one way to accommodate tasks is to cluster together states that frequently co-occur in the same task.

Casting hierarchy discovery as hidden state inference allows us to formalize this intuition with a straightforward extesion to our model. Following previous authors (Solway et al., 2014b; Balaguer et al., 2016), we assume the agent faces a sequence of tasks in *G*, where each task is to navigate from a starting state *s* ∈ *V* to a goal state *g* ∈ *V*. We assume the agent prefers shorter routes. Defining *tasks* = {*task_t_*} and *task_t_* = (*s_t_, g_t_*), we can extend the generative model with (Figure 3C):

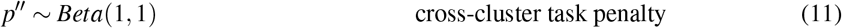

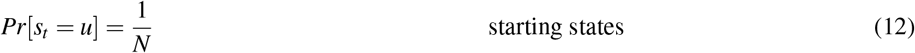

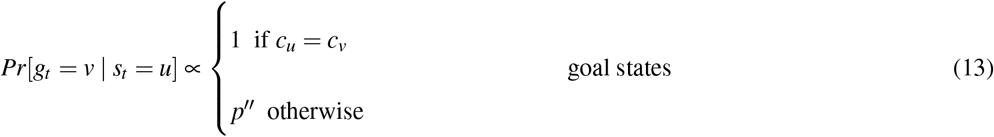

where *N* = |*V*| is the total number of states. Eq. 12 expresses the agent’s belief that a task can start randomly in any state. Eq. 13 expresses the belief that tasks are less likely to have goal states in a different cluster, with *p″* controlling exactly how much less likely that is.

If we denote the observable data as *D* = (*tasks, G*), the posterior becomes:

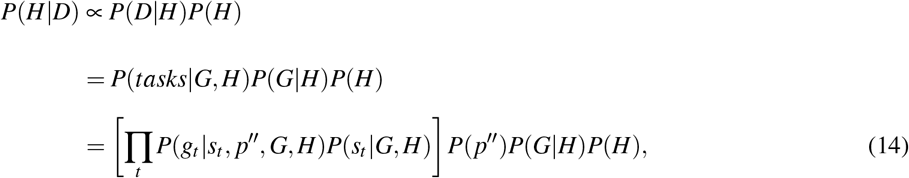

where the last two terms are the same as in Eq. 10.

The model will thus favor hierarchies that cluster together states which frequently co-occur in same tasks. This predicts that, in the absence of community structure, hierarchical planning will occur over clusters delineated by task start and goal states. This is a key novel prediction of our model which we assess in experiments one through five.

### Reward distribution

Besides topology and tasks, another factor that may play a role in hierarchy discovery is the distribution of rewards in the environment. In accordance with RL, we assume that each state delivers stochastic rewards and the agent aims to maximize the total reward (Sutton and Barto, 2018). Hierarchy discovery might account for rewards by clustering together states that deliver similar rewards. This is consistent with the tendency for humans to cluster based on perceptual features (Balaguer et al., 2016) and would be rational in an environment with autocorrelation in the reward structure (Srivastava et al., 2015; Schulz et al., 2018; Wu et al., 2018).

We can incorporate this intuition into the generative model by positing that states in the same cluster deliver similar rewards (Figure 3D):

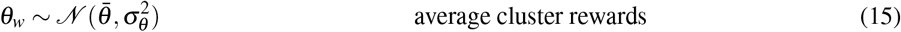

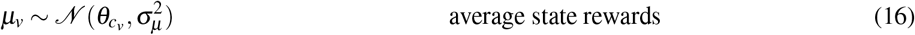

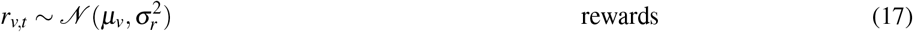

where *w* ∈ *V′* and *v* ∈ *V*. 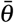 is the average reward of all states, *θ_w_* is the average reward of states in cluster *w, μ_v_* is the average reward of state *v, r_v,t_* is the actual reward delivered by state *v* at time *t*, and 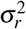 is the variance of that reward.

The observable data thus becomes *D* = (*r, tasks, G*) and the posterior can be computed as:

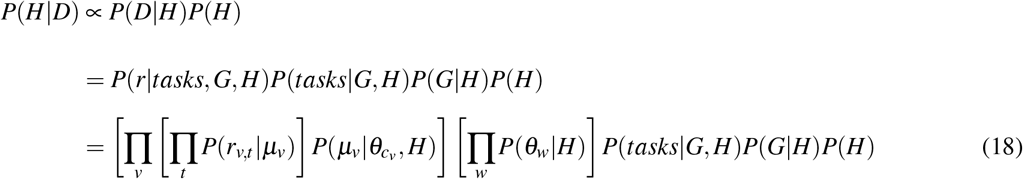

The model will thus favor hierarchies that cluster together states which deliver similar rewards. This predicts a certain pattern of reward generalization, with states inheriting the rewards of other states in the same cluster. Importantly, this implies that boundaries in the reward landscape will induce clusters that in turn will influence planning. This is another key novel prediction of our model which we assess in experiments six and seven.

### Inference

We approximated Bayesian inference using Markov chain Monte Carlo (MCMC) sampling (see Appendix) to draw samples approximately distributed according to *P*(*H*|*D*). We simulate each participant by drawing a single hierarchy *H* (the sampled hierarchy) from the posterior and then using it to make decisions. This is equivalent to assuming participants perform probability matching in the space of hierarchies.

In all simulations, we assume perfect (lossless) memory for the observations *D* = (*r, tasks, G*). While the process of learning the graph structure and the task and reward distributions is interesting in its own right, our focus is on inferring the hidden hierarchy *H*. We thus aim to develop a computational-level theory (in the Marrian sense; Marr and Poggio, 1976) of hierarchy discovery that remains agnostic of the particular algorithmic and implementational details but rather instantiates an entire class of process models that could approximate the ideal Bayesian observer.

### Choices

For simulations one through five, we simulate choices based on *H* using linking assumptions analogous to those used by the authors of the original studies. For experiments one through five and experiment seven, we simulate hierarchical planning based on *H* using HBFS. For experiment six, we assume participants prefer the state with the highest expected reward. For experiment eight, we assume participants prefer to reduce the entropy of the posterior as much as possible. In order to account for choice stochasticity, for each decision, we simulate the appropriate choice as dictated by the model with probability *ε*, or choose randomly with probability 1 – *ε*. We picked all model parameters by hand based on simulations one through five and based on the design for experiments six and seven. We used the same parameters throughout all simulations and experiments (Table 1).

**Table 1.** Model parameter settings. These were held constant across all simulations and experiments.

| Parameter | Range | Role | Value |
| --- | --- | --- | --- |
| $\alpha$ | $[0, +\infty)$ | CRP concentration parameter: larger values favor more clusters | 1 |
| $\varepsilon$ | $[0, 1]$ | choice stochasticity: larger values lead to more deterministic choices | 0.6 |
| # samples | $[1, +\infty)$ | number of MCMC iterations per simulated participant | 10000 |
| $\bar{\theta}$ | $(-\infty, +\infty)$ | average reward of entire graph | 15 |
| $\sigma_{\theta}$ | $(-\infty, +\infty)$ | standard deviation of average cluster rewards around $\bar{\theta}$ | 10 |
| $\sigma_{\mu}$ | $(-\infty, +\infty)$ | standard deviation of average state rewards | 10 |
| $\sigma_r$ | $(-\infty, +\infty)$ | standard deviation of state rewards | 5 |

## Simulations

### Simulation one: bottleneck transitions

Schapiro et al. (2013) demonstrated that people can detect transitions between states belonging to different clusters in a graph with a particular topological structure, such that nodes in certain parts of the graph are more densely connected with each other than with other nodes. This type of topology is also referred to as *community structure* (Figure 4A), with the clusters built into the graph topology referred to as *communities*. Thirty participants viewed sequences of stimuli representing random walks or Hamiltonian paths in the graph. Participants were instructed to press a key whenever they perceived a natural breaking point in the sequence. The authors found that participants pressed significantly more for cross-community transitions than for within-community transitions (Figure 4B). Participants did this despite never seeing a bird’s-eye view of the graph or receiving any hints of the community structure.

**Figure 4.**
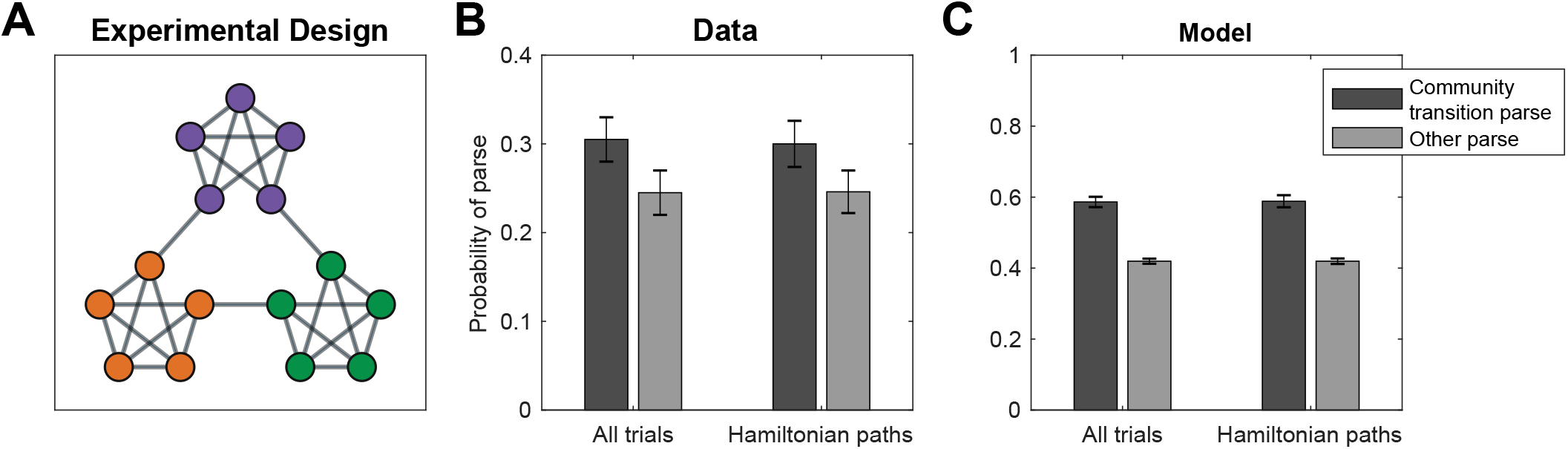
Detecting transitions between communities. A. Graph from Schapiro et al. (2013). Colors visualize the communities of states. Participants never saw the graph or received hints of the community structure. B. Results from Schapiro et al. (2013), experiment 1, showing that participants were more likely to parse the graph along community boundaries. Participants indicated transitions across communities as “natural breaking points” more often than transitions within communities. Error bars are s.e.m. (30 participants). C. Results from simulations showing that hierarchy inference using our model is also more likely to parse the graph along community boundaries. Error bars are s.e.m. (30 simulations).

Following experiment 1 from Schapiro et al. (2013), for each simulated participant, we sampled a hierarchy *H* based on the graph *G* and performed 18 random walks of length 15 initiated at random nodes, and 18 random Hamiltonian paths. For simplicity, we simulated key presses deterministically. In particular, we counted a transition from node *u* to node *v* as a natural breaking point if and only if the nodes belonged to different clusters in the inferred hierarchy *H*, that is, if *c_u_* ≠ *c_v_*, where *c* are the cluster assignments in *H*. This recapitulated the empirical results (Figure 4C; random walks: *t*(58) = 7.35, *p* < 10^−9^; Hamiltonian paths: *t*(58) = 7.32, *p* < 10^−9^, two sample, two-tailed *t*-tests). The inferred hierarchies for several simulated participants are shown in Figure 4D.

The dense connectivity within communities and sparse connectivity across communities drives the posterior to favor hierarchies with cluster assignments similar to the true underlying community structure. This arises due to Eq. 7 and Eq. 8 in the generative model, corresponding to desiderata 3 and 4, respectively. This posits that, during the generative process, edges across clusters are less likely than edges within clusters, resulting in a posterior distribution that penalizes alternative hierarchies in which many edges end up connecting nodes in different communities.

### Simulation two: bottleneck states

Solway et al. (2014a) performed several experiments demonstrating that people spontaneously discover graph decompositions that fulfill certain formal criteria of optimality. Similarly to Schapiro et al. (2013), in their first experiment, forty participants were trained on a graph with community structure (Figure 5A). As before, participants never saw the full graph or were made aware of its community structure but instead had to to rely solely on transitions between nodes. Participants were then asked to designate a single node in the graph as a “bus stop”, which they were told would reduce navigation costs in a subsequent part of the experiment. Participants preferentially picked the two bottleneck nodes on the edge that connects the two communities (Figure 5B), which are the optimal subgoal locations under these constraints. This suggests that participants were able to infer the graph topology based on adjacency information only, and to decompose it in an optimal way.

**Figure 5.**
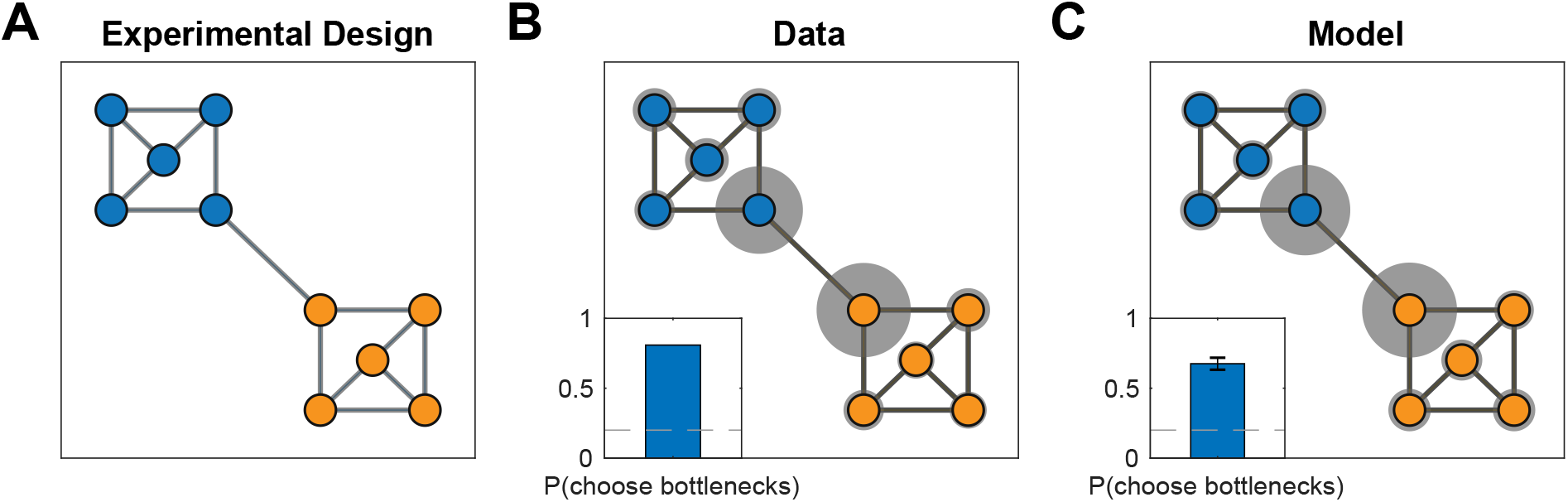
Detecting bottlenecks states. A. Graph from Solway et al. (2014a), experiment 1, with colors indicating the optimal decomposition according to their analysis. B. Results from Solway et al. (2014a), experiment 1, showing that people are more likely to select the bottleneck nodes as bus stop locations. Gray circles indicate the relative proportion of times the corresponding node was chosen. Inset, proportion of times either bottleneck node was chosen. Dashed line is chance (40 participants). C. Results from simulations showing that our model is also more likely to pick the bottleneck nodes since they are more likely to end up as endpoints of a bridge. Notation as in B. Inset error bars are s.e.m (40 simulations).

As in simulation one, for each participant we sampled a hierarchy *H* based on the graph *G* used in the experiment. Since participants were asked to identify three candidate bus stops, we randomly sampled three nodes among all nodes that belonged to bridges in *H*, i.e. {*u*: *b_w,z_* = (*u, v*), for some (*w,z*) ∈ *E′* and *u* ∈ *V*}, where the *b* and *E′* are the bridges and edges in the sampled hierarchy, respectively. This replicated the empirical result (Figure 5C; 65% of choices, *p* < 10^−20^, right-tailed binomial test), with most simulated participants inferring hierarchies respecting the underlying community structure (Figure 5D). Similarly to simulation one, this was due to the higher connectivity within communities than across communities.

### Simulation three: hierarchical planning

In their second experiment, Solway et al. (2014a) trained 10 participants to navigate between pairs of nodes in a different graph (Figure 6A). On some trials, participants were asked to indicate a single node that lies on the shortest path between two nodes in different communities. They found that participants overwhelmingly selected the bottleneck node between the two communities (Figure 6B), suggesting that they not only discovered the underlying community structure, but also leveraged that structure to plan hierarchically, “thinking first” of the high-level transitions between clusters of states.

**Figure 6.**
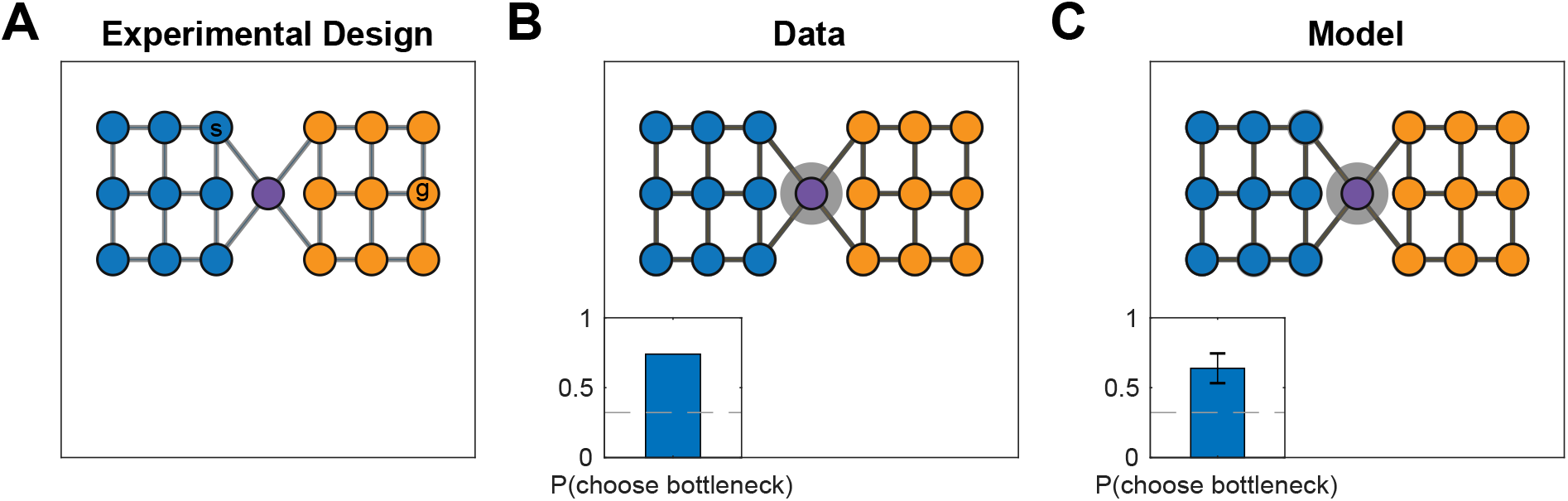
Planning transitions across communities first. A. Graph from Solway et al. (2014a), experiment 2, with colors indicating the optimal decomposition according to their analysis. The nodes labeled s and g indicate an example start node and goal node, respectively. B. Results from Solway et al. (2014a), experiment 2, showing that people are more likely to think of bottlenecks states first when they plan a path between states in different communities. Notation as in Figure 5B (10 participants). C. Results from our simulation demonstrating that our model also shows the same preference. Using the hierarchy identified by our model, the hierarchical planner is more likely to consider the bottleneck state first, since it is more likely to end up as the endpoint of a bridge connecting the two clusters. Error bars are s.e.m (10 simulations).

As in simulations one and two, we sampled *H* based on the graph *G* for each participant. We then used the hierarchical planner to find 50 hierarchical paths between random start locations in the left community and random goal locations in the right community. For each such path, we counted the first node that belongs to a bridge as the response on the corresponding trial, since this is the first node considered by the planner and is therefore the closest approximation to what a participant would think of first (see definition of HBFS in the Appendix). This replicated the empirical results, with a strong tendency for the bottleneck location to be selected (Figure 6C; 60% of choices, *p* < 0.001, right-tailed Monte Carlo test). The discovered hierarchies resembled the underlying community structure for the same reason as in simulations one and two (Figure 6D), resulting in the bottleneck node frequently becoming part of a bridge that all paths between the two communities would pass through.

### Simulation four: shorter hierarchical paths

In their final experiment, Solway et al. (2014a) demonstrated hierarchical decomposition and planning in the Towers of Hanoi task, which can be represented by a graph (Figure 7A) in which each node is a particular game state and each edge corresponds to move that transitions from one game state to another. They leveraged the fact that there are two different shortest paths between some pairs of states (for example, the start and goal states in Figure 7A), but those paths cross a different number of community boundaries as defined by their optimal decomposition analysis. Hierarchical planning predicts that participants will prefer the path which crosses fewer community boundaries, and this is indeed what the authors found (Figure 7B).

**Figure 7.**
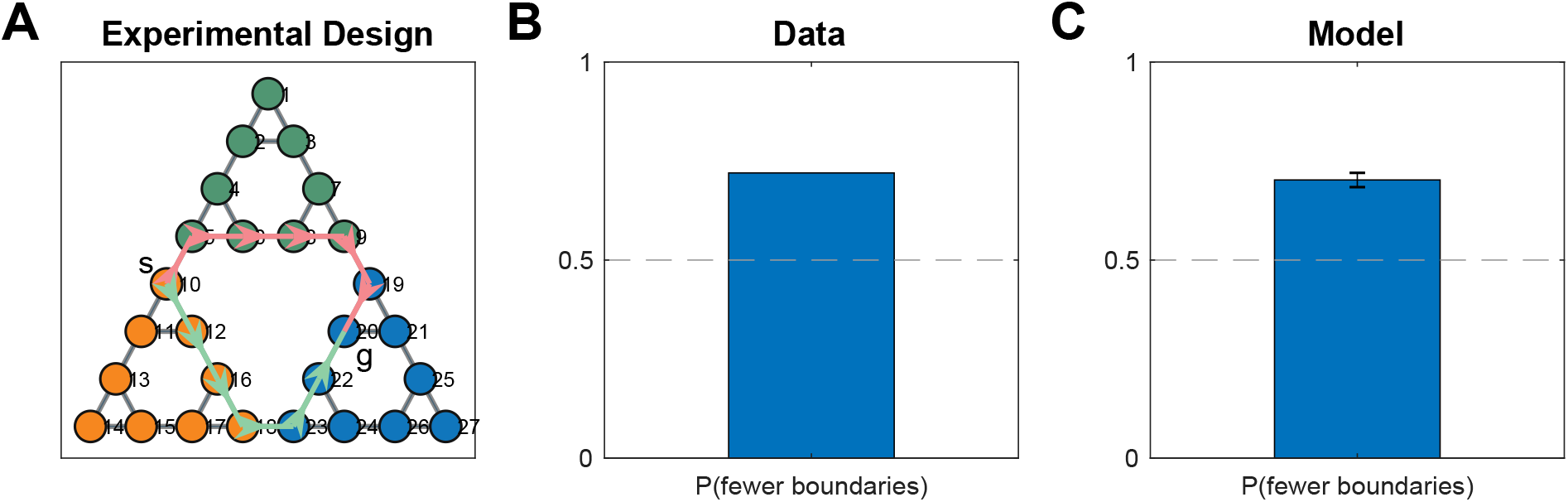
Preferring paths with fewer community boundaries. A. Graph representing the Towers of Hanoi task used in Solway et al. (2014a), experiment 4. Vertices represent game states, edges represent moves that transition between game states. The start and goal states (*s, g*) show an example of the kinds of tasks used in the experiment. Colored arrows denote the two shortest paths that could accomplish the given task, with the red path passing through two community boundaries and the green path passing through a single community boundary. B. Results from Solway et al. (2014a), experiment 4, showing that participants preferred the path with fewer communities, or equivalently, the path that crosses fewer community boundaries. Bar graph shows fraction of participants (35 participants). Dashed line is chance. C. Results from simulations showing that our model also exhibits the same preference. Bar graph shows the fraction of simulations that chose the path with fewer community boundaries. Error bar is s.e.m. (35 simulations).

After choosing a hierarchy *H* for each simulated participant as in the previous simulations, we then used the hierarchical planner to find paths between the six pairs of states that satisfy the desired criteria. In accordance with the data, this resulted in the path with fewer community boundaries being selected more frequently (Figure 7C; 71.4% of choices, *p* < 10^−10^, right-tailed binomial test). Similarly to the previous simulations, the model tended to carve up the graph along community boundaries (Figure 7D). Since the planner first plans in the high-level graph, it prefers the path with the fewest clusters, and hence with the fewest cluster boundaries.

### Simulation five: cross-cluster jumps

In our final simulation, we considered a study performed by Lynn et al. (2018), in which participants experienced a random walk along the graph shown in Figure 8A and had to respond to each stimulus (node) after each transition. A small subset of transitions violated the graph structure and instead “teleported” the participant to a node that is not connected to the current node. Importantly, there were two types of violations: short violations of topological distance 2 and long violations of topological distance 3 or 4. The authors found that participants were slower to respond to longer than to shorter violations, suggesting that participants had inferred the large scale structure of the graph.

**Figure 8.**
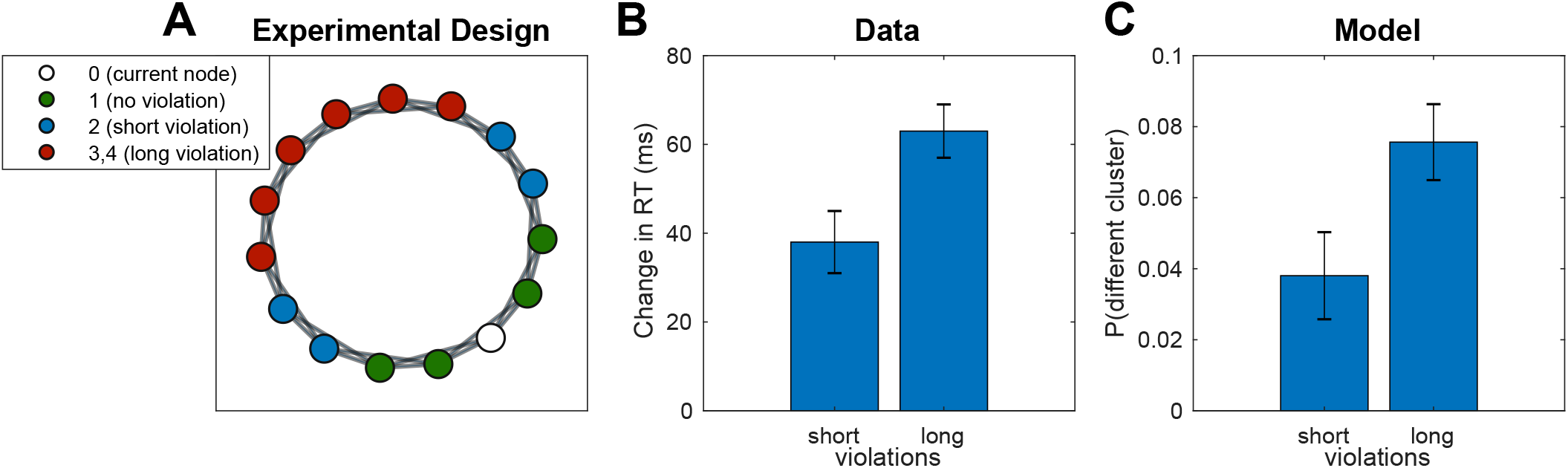
Slower reactions to cross-cluster transitions. A. Graph used in Lynn et al. (2018). Each node (white) is connected to its neighboring nodes and their neighbors (green). Blue nodes are 2 transitions away from the white node, while red nodes are 3 or 4 transitions away. B. Results from Lynn et al. (2018) showing that, on the test trial, participants were more slower to respond to long violations than to slow violations. Change in RT is computed with respect to average RT for no-violation transitions. Error bars are s.e.m (78 participants). RT, reaction time. C. Results from simulations showing that long violations are more likely to end up in a different cluster, which would elicit a greater surprise and hence a slower RT, similar to crossing a cluster boundary.

Like in the previous simulations, we sampled *H* for each participant and then simulated a random walk along the graph G, with occasional violations as described in Lynn et al. (2018). In order to model reaction times (RT’s), we assumed a bimodal distribution of RT’s, with fast RT’s for transitions within clusters and slow RT’s for transitions across clusters, consistent with the notion that cross-cluster transitions are more surprising. Instead of actually simulating RT’s, we simply counted the number of cross-cluster transitions during the random walk. This revealed a greater number of cross-cluster transitions for long violations than for short violations, consistent with the data (Figure 8B; *t*(154) = 4.50, *p* = 0.00001, two-sample t-test). This occurs because nearby nodes are more likely to be clustered together (Figure 8D), and hence violations of greater topological distance increase the likelihood that the destination node would be in a different cluster than the starting node, resulting in a greater surprise and a slower RT.

## Experiments

### Experiment one: task distributions

In our first experiment, we sought to validate the prediction of our model that clusters could be induced by the distribution of tasks alone, even when the graph topology does not favor any particular clustering. In particular, our model predicts that states which frequently co-occur in the same task should be clustered together, since the hierarchical planner is optimal within clusters.

#### Participants

We recruited 87 participants (28 female) from Amazon Mechanical Turk (MTurk). All participants received informed consent and were paid $3 for their participation. We reasoned that paying participants a fixed amount would incentivize them to complete the experiment in the least amount of time possible, which entails balancing path length with planning time in a way that is characteristic of many real-world tasks. The experiment took 17 minutes on average. All experiments were approved by the Harvard Institutional Review Board.

#### Design

We asked participants to navigate between pairs of nodes (“subway stops”) in a 10-node graph (the “subway network”, Figure 9A, left). The training trials (Figure 9A, right) were designed to promote a particular hierarchy: the task to navigate from node 1 node 3 would favor clustering nodes 1,2,3 together; the task to navigate from node 4 to node 6 would favor clustering nodes 4,5,6 together; and the task to navigate from node 10 to node 7 would favor clustering nodes 7,8,9,10 together (Figure 9A, left). The normative reason for this is that hierarchical planning is always optimal within a cluster. In the generative model, this is taken into account by Eq. 13, which leads to a preference to cluster together start and goal states from the same task. The purpose of the tasks with random start and goal states was to encourage participants to learn a state representation for efficient planning and not simply to respond habitually.

**Figure 9.**
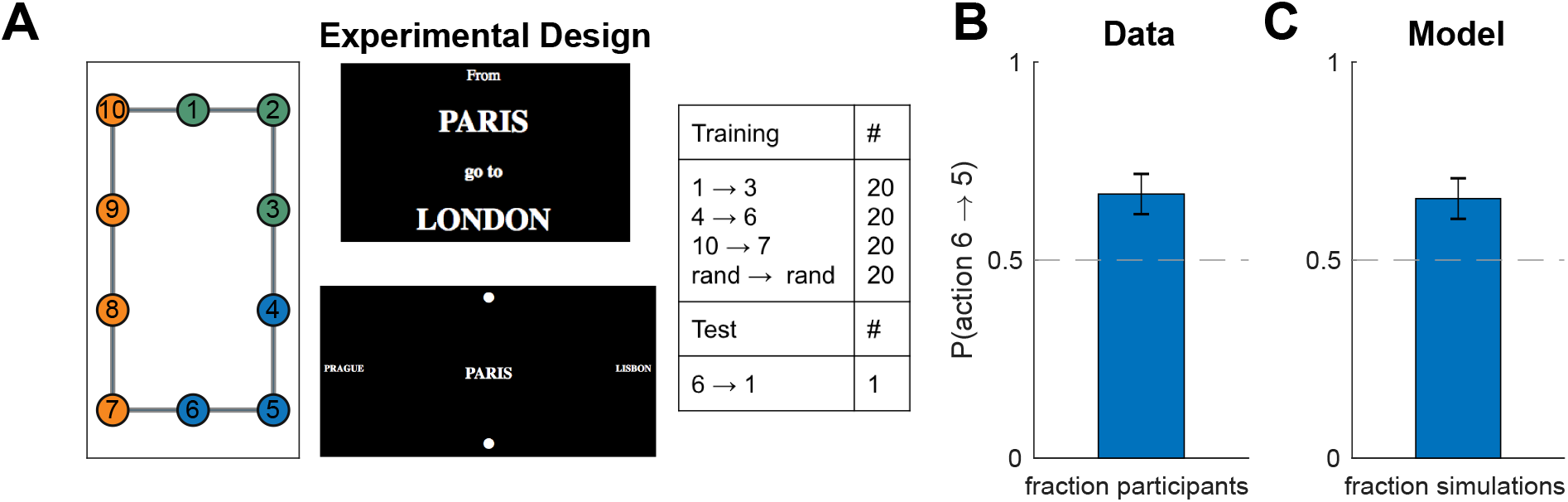
Hierarchy discovery is sensitive to the task distribution. A. (Left) graph used in experiment one with no topological community structure. Colors represent clusters favored by the training protocol (right). Numbers serve as node identifiers and were not shown to participants. “Rand” denotes a node that is randomly chosen on each trial. (Middle) trial instruction (top) and screenshot from the starting state (bottom). B. Results from experiment one showing that, on the test trial, participants were more likely to go to state 5 than to state 7, indicating a preference for the route with fewer cluster boundaries. Dashed line is chance. Error bars are s.e.m. (87 participants) C. Results from simulations showing that our model also preferred the transition to state 5. Notation as in B.

In order to test the model prediction, after training, we asked participants to navigate from node 6 node 1. Note that the two possible paths are of the same length and a planner with a flat representation of the graph would show no preference for one path over the other. Furthermore, since there is no community structure and the graph is perfectly symmetric, any clustering strategy based on graph structure alone would not predict a preference. Conversely, our model predicts that participants will tend to choose the path through node 5 since it passes through a single cluster boundary, whereas the path through node 7 passes through two cluster boundaries.

#### Procedure

The experiment was implemented as a computer-based game similar to Balaguer et al. (2016) in which participants had to navigate a virtual subway network. At the start of each trial, participants saw the names of the starting station and the goal station. After 2 s, they transitioned to the navigation phase of the trial, during which they could see the name of the current station in the middle of the screen, surrounded by the names of the four neighboring stations, one in each cardinal direction. If there was no neighboring station in a particular direction, participants saw a filled circle instead of a station name. The name of the goal station was also indicated in the top left corner of the screen as a reminder. The navigation phase began with a 3-s countdown during which participants could see the starting station and its neighbors but could not navigate. Participants were instructed to plan their route during the countdown. After the countdown, participants could navigate the subway network using the arrow keys. Transitions between stations were instantaneous. Once participants reached the goal station, they had to press the space bar to complete the trial. This was followed by a 500-ms “success” message, after which the trial ended and the instruction screen for the next trial appeared. Pressing the space bar on a non-goal station resulted in a “incorrect” message flashing on the screen. Attempting to move in a direction without a neighboring station had no effect. Following Balaguer et al. (2016), stations were named after cities, with the names randomly shuffled for each participant.

The subway network corresponded to the graph in Figure 9A. In order to assign arrow keys to edges, we first embedded the graph in the Cartesian plane by assigning coordinates to each vertex, which resulted in the planar graph shown in the figure. Then we assigned the arrow keys to the corresponding cardinal directions. For each participant, we also randomly rotated the graph by 0°, 90°, 180°, or 270°. Participants performed 80 training trials (20 in each condition; Figure 9, right) in a random order. After the training trials, we showed a message saying that now the subway system was unreliable, so that some trips may randomly be interrupted midway. This was followed by the test phase, during which participants performed the test trial 6 → 1 and two additional test trials. In order to prevent new learning during the test phase, all test trials were interrupted immediately after the first valid keypress. The two additional test trials were not included in the analysis or any of the following experiments. We used the destination of the first transition on the test trial as our dependent measure in the analysis. We reasoned that the direction in which participants attempt to move first is along what they perceive to be the best route to the goal station. Since participants were paid a fixed fee for the whole experiment, they were incentivized to complete it as fast as possible, which can be best achieved by planning the shortest route during the 3-s countdown and then following it.

#### Results and discussion

In accordance with our model predictions, more participants moved to state 5 on the test trial, rather than to state 7 (Figure 9B; 58 out of 87, *p* = 0.003, two-tailed binomial test). Notice that this could not be explained by habitual responding: while participants may have learned action chunks that solve the corresponding tasks (for example, pressing right and down from state 1 to state 3), the actions from state 6 to its neighboring states were never reinforced as part of a stimulus-response association or as part of a longer action chunk. In particular, state 6 is never a starting state or an intermediary state, except possibly in the random tasks, in which both directions have equal probability of being reinforced. The effect cannot be explained by state familiarity either, since participants experienced states 5 and 7 equally often, on average. Finally, it is worth noting that a model-free RL account would make the opposite prediction, since state 7 would on average have a higher value than state 5, as it gets rewarded directly.

This result was consistent with the model predictions (Figure 9C; 57 out of 87, *p* = 0.005, two-tailed binomial test), suggesting that people form hierarchical representations for planning in a way that tends to cluster together nodes that co-occur in the same tasks.

### Experiment two: task distributions and suboptimal planning

Since both paths on the test trial in the previous experiment were of equal length, the bias that participants developed would make no difference for their performance on that task. Next we asked whether such a bias would occur even if it might lead to suboptimal planning, as the hierarchical planner would predict. For example, most of us have had the experience of navigating between two locations through other, more familiar location, even though a shorter but less familiar route might exist (see also Huys et al., 2015).

#### Participants

We recruited 241 participants (112 female) from MTurk. Of those, 78 were assigned to the “bad” clusters condition, 87 were assigned to the “control” condition, and 76 were assigned to the “good” clusters condition. Participants were paid $2.50 for their participation. The experiment took 15 minutes on average.

#### Design and procedure

We used the same paradigm as experiment one, with the only difference that node 10 was removed from the graph (Figure 10A, left). Now the analogous training regime (Figure 10A, right) would promote “bad” clusters that lead to suboptimal planning on the test trial. We also performed a control version of the experiment with different participants using random training tasks only, as well as a “good” condition with a third group of participants that promotes clusters that lead to optimal planning on the test trial (Figure 10A).

**Figure 10.**
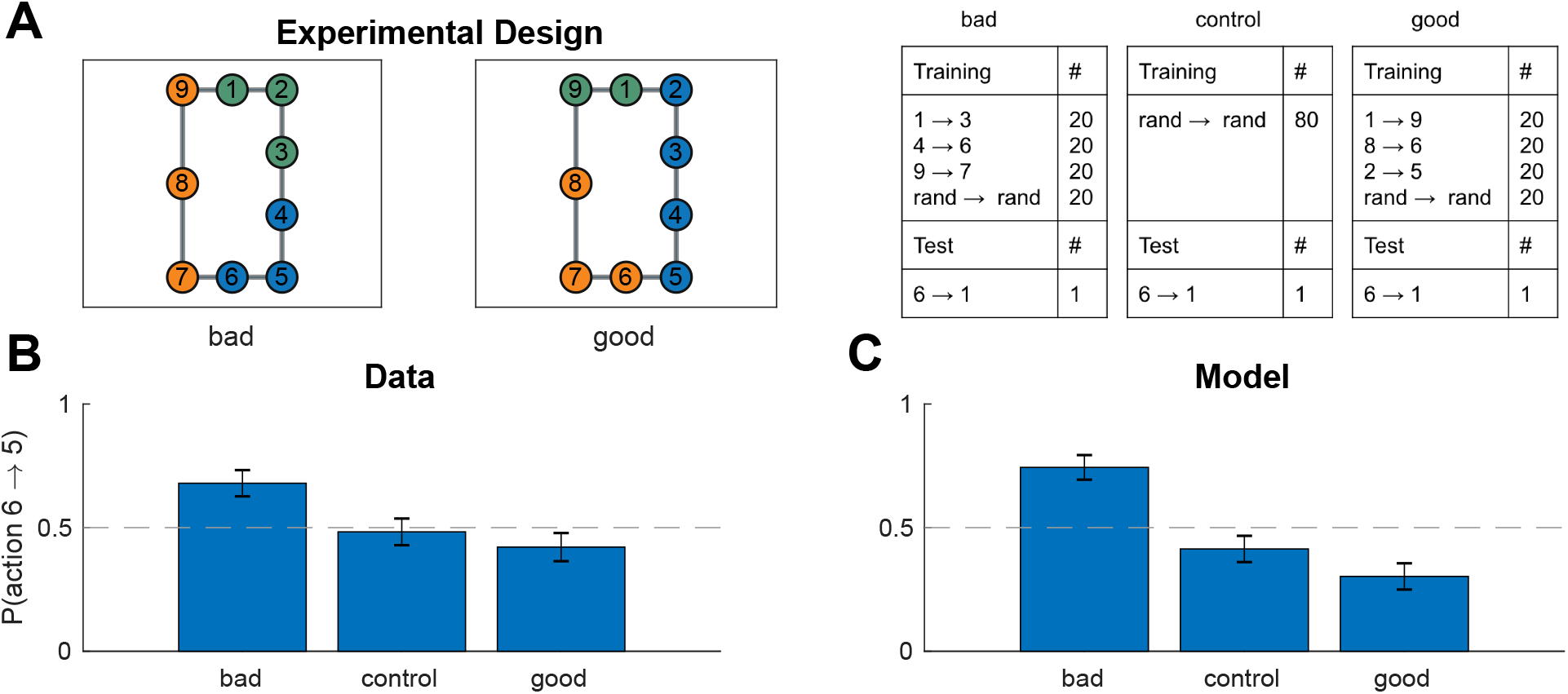
Different task distributions can induce different hierarchies in the same graph. A. (Left) graph used in experiment two with colors representing clusters favored by the training protocol in the “bad” (left) and “good” (middle) condition. (Right) training and test protocols for all three conditions. B. Results from experiment two showing that, on the test trial, participants were more likely to go to state 5 than to state 7 in the bad condition, leading to the suboptimal route. The effect was not present in the control condition or in the good condition. Dashed line is chance. Error bars are s.e.m. (78, 87, and 76 participants, respectively). C. Results from simulations showing that our model exhibited the same pattern. Notation as in B.

#### Results and discussion

On the test trial, participants preferred the suboptimal move from 6 to 5 in the “bad” clusters condition (Figure 10B; 53 out of 78, *p* = 0.002, two-tailed binomial test), significantly more than the control condition (*χ*^2^(1,165) = 6.52, *p* = 0.01, chi-square test of independence) and the “good” condition (*χ*^2^(1,154) = 10.4, *p* = 0.001). This was in accordance with the model predictions (Figure 10C; 58 out of 78 simulated participants, *p* = 0.00002, in the bad condition; *χ*^2^(1,165) = 18.2, *p* = 0.00002 for bad vs. control condition; *χ*^2^(1,154) = 30.0, *p* < 10^−7^ for bad vs. good condition). This suggests that participants formed clusters based on the distribution of tasks and used these clusters for hierarchical planning, even when that was suboptimal on the particular test task. Somewhat surprisingly, there was no difference between the control condition and the good condition in the data (*χ*^2^(1,163) = 0.6, *p* = 0.5), and the model showed the same pattern (*χ*^2^(1,163) = 2.2, *p* = 0.14), suggesting that in this particular scenario the “good” clusters have a relatively weaker effect.

### Experiment three: learning effects

While the experiments so far demonstrate the key predictions of our model, they only test asymptotic behavior and as such do not assess how beliefs about hierarchy evolve over the course of learning. Further, these results could conceivably be explained by simpler, non-hierarchical accounts, such as a bidirectional association forming between states 5 and 6 due to the frequent transition from 5 to 6 during training. The goal of our next experiment was to rule out such “flat” associative explanations and to study the dynamics of learning.

#### Participants

We recruited 127 participants (54 female) from MTurk. Participants were paid $8.50 for their participation. The experiment took 47 minutes on average.

#### Design

We trained participants on the same graph experiment one, but using a different training regimen, with two stages of training and multiple probe trials (Figure 11A). The first stage of training (trials 1-103) promoted clustering states 2 and 3 together, separately from state 1, and similarly promoted clustering states 4 and 5 together, separately from state 6 (Figure 11A, first panel). In order to investigate the dynamics of hierarchy discovery, we interspersed three probe trials that tasked participants to navigate from state 6 to state 1 throughout the first stage, expecting participants’ preferences to become stronger over time. Note that predictions on the probe trials are the opposite of predictions on the test trial in experiment one: the path from 6 to 1 through state 5 now crosses three cluster boundaries, whereas the path through state 7 crosses only two cluster boundaries, so participants should prefer the path through state 7. This cannot be explained by a naive associative account since state 6 is no more likely to co-occur with state 7 than with state 5.

**Figure 11.**
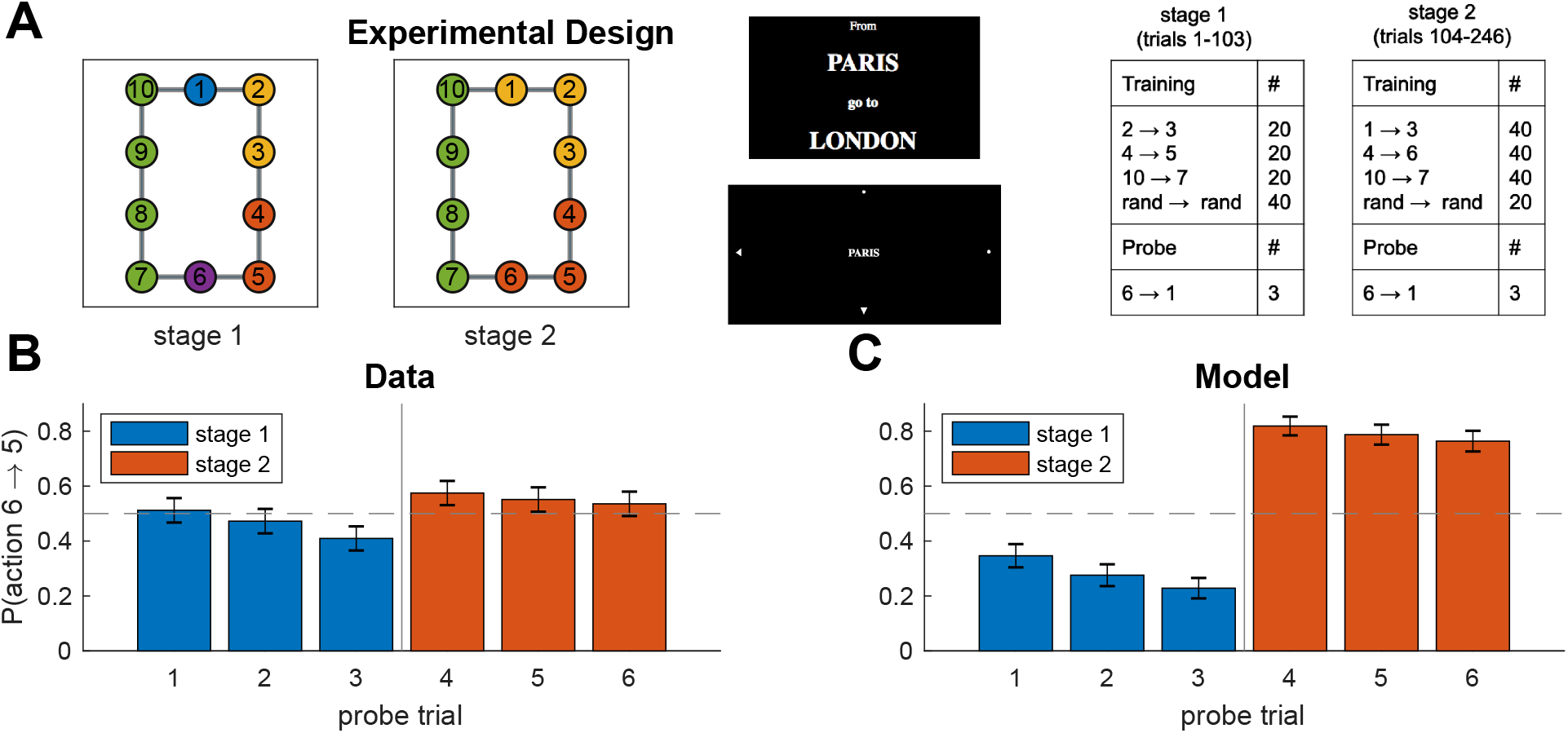
Learning dynamics. A. Experiment three used the same graph as experiment one, with main difference that training (right panel) took part in two stages that promoted different hierarchies (first and second panel), with probe trials interspersed throughout training. Notation as in Figure 9A. B. Results from experiment three showing that (1) the first stage of training makes participants more likely to go to state 7 on the probe trials, which could not be explained by a “flat” associative account, (2) this tendency appears gradually as participants accumulate more evidence, and (3) this preference is reversed during the second stage of training. Error bars are s.e.m. (127 participants). C. Results from simulations showing that our model exhibited the same learning dynamics. Notation as in B.

To further assess learning, which sought to reverse this effect during the second stage of training which promoted the same clusters as experiment one (Figure 11A, second panel). Our prediction was that, in accordance with the results of experiment one, this would eliminate participants’ preference for going to state 7 on the probe trials in favor of state 5, since the path through state 5 now crosses a single cluster boundary.

#### Procedure

We used the same procedure as experiment one, with the following changes. First, there was no information indicating to participants that something had changed between stages (i.e., between trials 103 and 104). Additionally, instead of having a test stage in the end, we interspersed six probe trials 6 → 1, spaced evenly throughout training (trials 34, 68, 103 during the first stage and trials 150, 197, 246 during the second stage). Unlike the test trials in experiment one, probe trials were indistinguishable from training trials and were not interrupted after the first move. Another difference from experiment one was that instead of being rotated, the graph was randomly flipped horizontally and/or vertically for each participant. Finally, unlike experiment one, participants did not see the names of adjacent stations but instead saw arrowheads indicating they could move in the corresponding direction (Figure 11A, third panel).

#### Results and discussion

Consistent with our predictions, participants showed a significant preference for moving to state 7 on the last probe trial of the first stage (Figure 11B, probe trial 3; 75 out of 127 participants, *p* = 0.05, two-tailed binomial test). This effect was modulated by the amount of training, with the smallest effect on the first probe trial and the largest effect on the third probe trial (slope = −0.27, *F*(1,379) = 3.96, *p* = 0.05, mixed effects logistic regression with probe trials 1-3). The effect was reversed during the second stage (slope = 0.73, *F*(1,252) = 8.10, *p* = 0.005, mixed effects logistic regression with probe trials 3-4).

These results were consistent with the model predictions (Figure 11C). The model preference for state 7 on the third probe trial (98 out of 127 simulated participants, *p* < 10^−9^, two-tailed binomial test) is once again due to the preference of the hierarchical planner for paths with fewer state clusters. As in our empirical data, this effect became stronger over time (slope = −0.41, *F*(1,379) = 6.82, *p* = 0.009, mixed effects logistic regression with probe trials 1-3). The reason is that, as the hierarchy inference process observes more tasks, it accumulates more evidence (i.e., more terms in the product in Eq. 14) which sharpens the posterior distribution *P*(*H*|*D*) around its mode (i.e., the most probable hierarchy, Figure 11A, first panel). This corresponds to a decrease in the uncertainty over *H*, which makes it more likely that the sampler will draw the hierarchy in Figure 11A, first panel, and plan according to it. Hence this effect of learning occurs due to the decrease in uncertainty resulting from the additional observations. Finally, like our participants, the model reversed its preference during the second stage of training (slope = 2.73, *F*(1,252) = 76.03, *p* < 10^−10^, mixed effects logistic regression with probe trials 3-4). This effect occurs because tasks during the second stage of training shift the mode of the posterior (Figure 11A, second panel). Together, these results rule out simple, nonhierarchical associative accounts and demonstrate that our model can account for learning dynamics due to changes in uncertainty and changes in the mode of the posterior distribution over hierarchies.

### Experiment four: perfect information

One downside of experiments one and two is that effects of memory confound hierarchy inference and planning. In particular, it is unclear that participants are able to learn and represent the full graph *G*. This is most evident in the control condition of experiment two (Figure 10B), in which our model predicts that participants should be better than chance, when in fact they are not, thus questioning whether they are learning the graph or planning efficiently in the first place. Note that this does not pose a significant challenge to our hierarchy discovery account; people’s choices could still be accounted for without invoking a hierarchical planner, for example by assuming a preference to remain in the same cluster (moving to state 5) rather than crossing a cluster boundary (moving to state 7). Nevertheless, we sought to overcome this limitation by ensuring participants know the full graph *G*.

#### Participants

We recruited 77 participants (33 female) from MTurk. Participants were paid $2.00 for their participation. The experiment took 10 minutes on average.

#### Design

We used the same graph and training protocol as experiment one, except this time participants could see the whole graph at any given time (Figure 12A). This put participants on an equal footing with the hierarchy inference and hierarchy planning algorithms, both of which assume perfect knowledge of the graph *G*.

**Figure 12.**
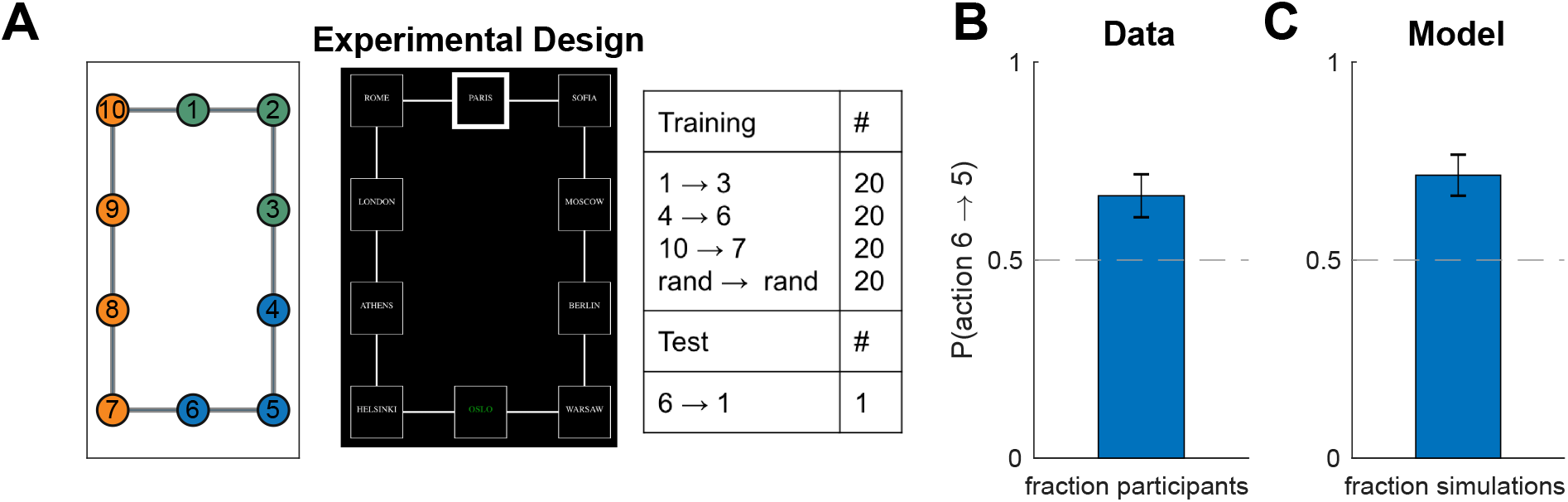
Hierarchy discovery based on task distribution in fully visible graphs. A. (Left) experiment four used the same graph as experiment one, however this time the graph was fully visible on each trial (middle). Notation as in Figure 9A. B. Results from experiment four showing that, like in experiment one, participants were more likely to go to state 5 on the test trial. Dashed line is chance. Error bars are s.e.m. (77 participants) C. Results from simulations showing that our model also preferred the transition to state 5. Notation as in B.

#### Procedure

The procedure was similar to experiment one, with the main difference that participants had a bird’s-eye view of the entire subway network throughout the experiment (Figure 12A, middle panel). Subway stations were represented by squares connected by lines which represented the connections between stations. The current station was highlighted with a thick border and the goal station was in green font.

Since planning is significantly easier in the setting, we removed the 3-s countdown, so that participants could start navigating immediately after the 2-s instruction. Additionally, instead of rotating the map, we randomly flipped it horizontally for half of the participants. We also omitted the “unreliable trips” warning before the test phase since participants now only saw a single test trial, which was immediately interrupted after the first move. The experiment took 10 minutes on average and participants were paid $2.00.

#### Results and discussion

We found that, even with full knowledge of the graph, participants still developed a bias (Figure 12B; 51 out of 77 participants, *p* = 0.0014, right-tailed binomial test), consistent with the predictions of our model (Figure 12C; 55 out of 77, *p* = 0.0002, two-tailed binomial test). This provides strong support for our hierarchy discovery account, suggesting tasks constrain cluster inferences above and beyond constraints imposed by graph topology, which in turn constrain hierarchical planning on novel tasks.

### Experiment five: perfect information and suboptimal planning

We next asked whether hierarchy discovery could lead to suboptimal planning even when the full graph is visible.

#### Participants

We recruited 386 participants (175 female) from MTurk. Of those, 119 were assigned to the “bad” clusters condition, 90 were assigned to the “control 1” condition, 88 were assigned to the “control 2” condition, and 89 were assigned to the “good” clusters condition. Participants were paid $2.00 for their participation. The experiment took 9 minutes on average.

#### Design and procedure

We used the same graph and training protocol as experiment two, except this time participants could see the whole graph at any given time (Figure 13A), as in experiment four. Additionally, we included two control conditions, one with 20 training trials (“control 1”) and one with 80 training trials (“control 2”). The first control condition ensured participants received the same number of random tasks as the bad and good conditions, while the second control condition ensured that participants received the same total number of tasks as in the bad and good conditions. We used the same experimental procedure as experiment four.

**Figure 13.**
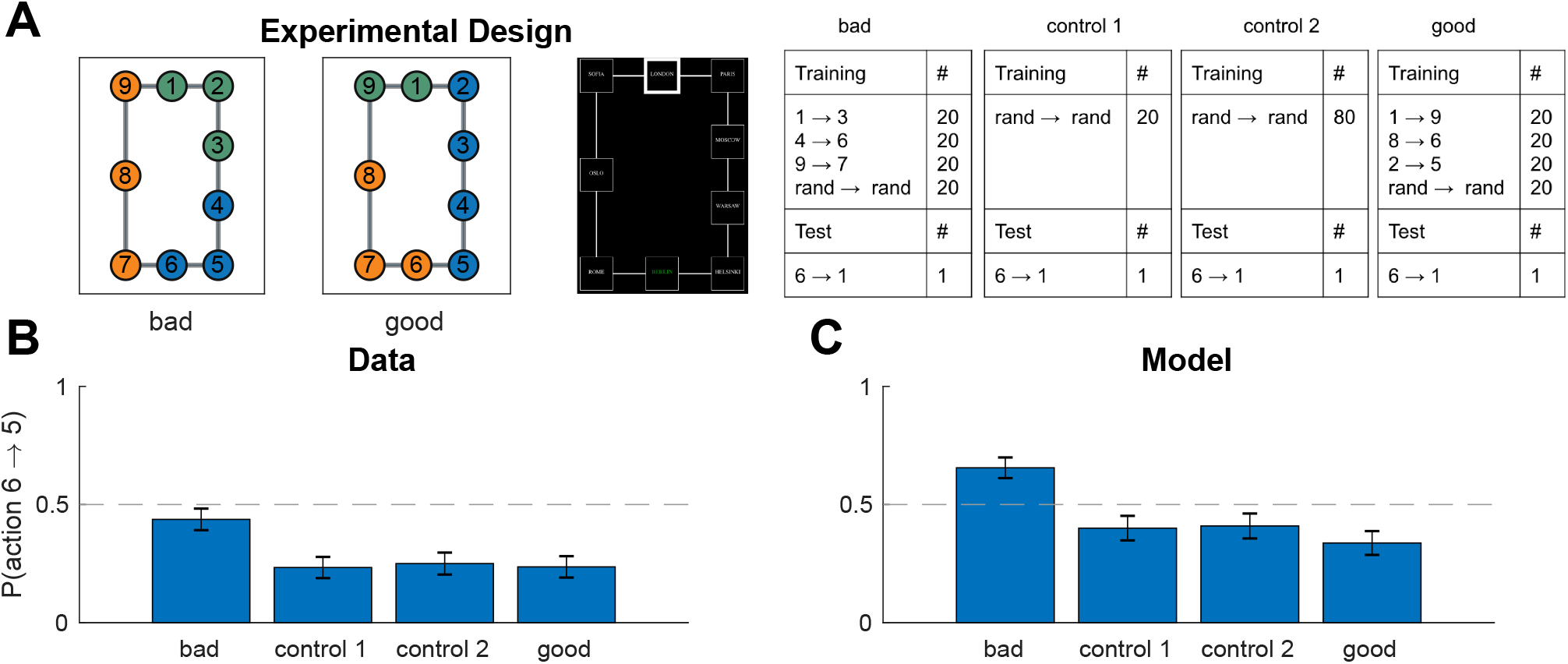
Task distributions can bias hierarchical planning even in fully visible graphs. A. (Left) experiment five used the same graph as experiment two, however this time the graph was fully visible on each trial. Notation as in Figure 10A. B. Results from experiment five showing that participants were still biased by the training tasks in the bad condition, performing worse on the test trial compared to the other conditions. Dashed line is chance. Error bars are s.e.m. (119, 90, 88, and 89 participants, respectively). C. Results from simulations showing that our model exhibited the same pattern. Notation as in B.

#### Results and discussion

As in experiment two, inducing “bad” clusters still led to significantly worse performance on the test trial than either control condition (Figure 13B; *χ*^2^(1,209) = 9.35, *p* = 0.002 for bad vs. control 1; *χ*^2^ (1,207) = 7.7, *p* = 0.006 for bad vs. control 2, chi-square test of independence). Inducing “good” clusters (89 participants) led to significantly better performance than “bad” clusters (*χ*^2^(1,208) = 9.03, *p* = 0.003 for bad vs. good), although not significantly better than the control conditions (*χ*^2^(1,179) = 0.002, *p* = 0.96 and *χ*^2^(1,177) = 0.05, *p* = 0.8 for good vs. control 1 and good vs. control 2, respectively). This accords with our model predictions (Figure 13C; *χ*^2^(1,209) = 10.1, *p* = 0.002 for bad vs. control 1; *χ*^2^(1,207) = 4.8, *p* = 0.03 for bad vs. control 2; *χ*^2^(1,208) = 17.7, *p* = 0.00003 for bad vs. good) and strongly suggests that people default to hierarchical planning over clusters influenced by the task distribution, even in simple, fully observable graphs. Notice that in both control conditions, participant preferred the shorter path (21 out of 90 participants, *p* < 10^−6^ for control 1; 22 out of 88 participants, *p* < 10^−6^ for control 2, two-tailed binomial tests), indicating that they were indeed able to plan effectively when given the full graph without tasks to bias them towards particular clusters, thus overcoming the limitation of experiment two.

One notable difference between our model predictions and the empirical data is that our model predicts a preference for state 5 in the “bad” condition (78 out of 119 simulated participants, *p* = 0.0009, two-tailed binomial test), whereas participants did not show significant preference (52 out of 119 participants, *p* = 0.2, two-tailed binomial test). We believe this occurs because the task is much simpler when the graph is fully visible and participants could easily perform optimal “flat” planning, rather than having to resort to hierarchical planning. This effect could be captured straightforwardly using a mixture of BFS and HBFS for planning, rather than just HBFS.

### Experiment six: reward generalization

In this experiment, we tested the prediction that rewards generalize within clusters. While this prediction is not unique to our model and could be accounted for by Gaussian processes over graphs (Kondor and Lafferty, 2002; Wu et al., 2018) or by the successor representation (Stachenfeld et al., 2017; Dayan, 1993), the idea that clusters generate similar rewards is a core assumption of our model that we sought to validate before assessing how clusters inferred based on rewards could influence planning (experiment seven).

#### Participants

We recruited 32 participants from the MIT undergraduate community. The experiment took around 3 minutes and participants were not paid for their participation.

#### Design

We showed participants the graph (“network of gold mines”) in Figure 14A and told them that in the past, states delivered an average reward of 15 (“grams of gold”), but today, state 4 (green) delivered a reward of 30. We then asked participants to choose one between state 3 and state 7 (“mines to explore”).

**Figure 14.**
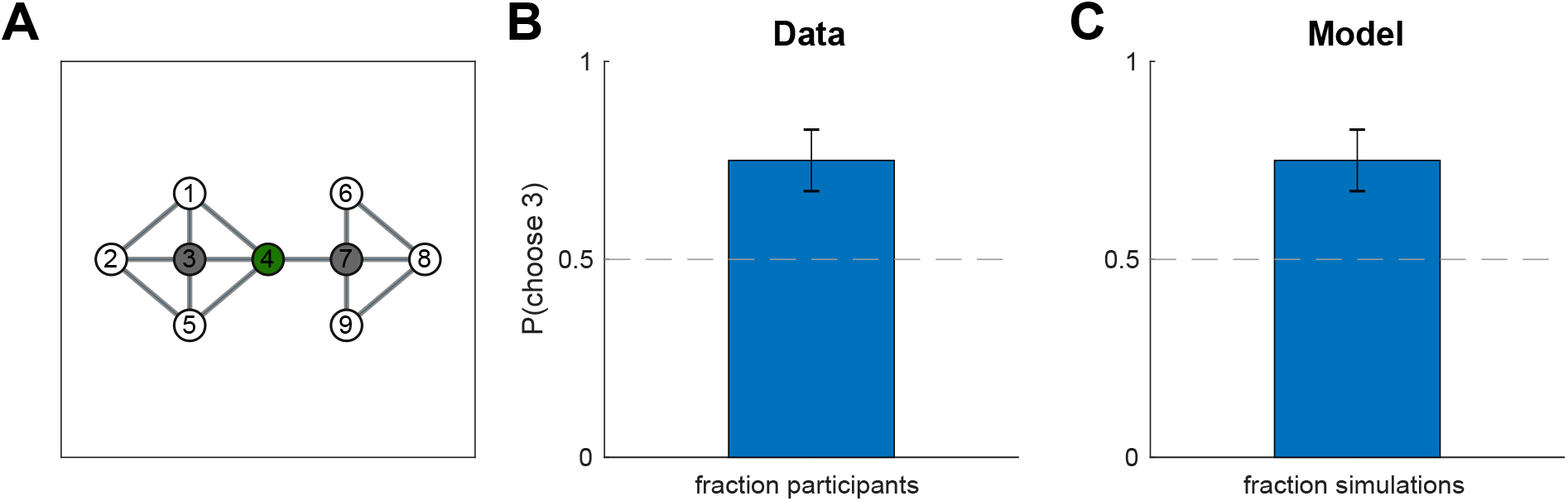
Reward generalization within clusters. A. Graph used in experiment six. Numbers indicate state identifiers and were not shown to participants. Participants were told that states deliver 15 points on average and that, on a given day, state 4 (green) delivered 30 points. They were then asked which of the two gray nodes (states 3 and 7) they would choose. B. Results from experiment six showing that participants preferred state 3, which is in the same topological cluster as state 4, suggesting they generalized the reward within the cluster. Error bars are s.e.m (32 participants). C. Results showing that the model exhibited the same pattern. Notation as in B.

#### Procedure

Each participant was given a sheet of paper with instructions and the graph in Figure 14A, without node identifiers. The instructions were as follows:

*You work in a large gold mine that is composed of multiple individual mines and tunnels. The layout of the mines is shown in the diagram below (each circle represents a mine, and each line represents a tunnel). You are paid daily, and are paid $10 per gram of gold you found that day. You dig in exactly one mine per day, and record the amount of gold (in grams) that mine yielded that day. Over the last few months, you have discovered that, on average, each mine yields about 15 grams of gold per day. Yesterday, you dug in the blue mine in the diagram below, and got 30 grams of gold. Which of the two shaded mines will you dig in today? Please circle the mine you choose*.

Half of the participants were given a version in which the graph was flipped horizontally, i.e. the topological cluster was on the right side.

#### Results and discussion

Participants preferred state 3, the state in the same topological community as state 4 (Figure 14B; 24 out of 32 participants, *p* = 0.007, two-tailed binomial test), as the model predicts (Figure 14C; 24 out of 32 simulated participants, *p* = 0.007). Topological structure is the only driver of hierarchy discovery in this case, since there is only a single reward and no tasks. The structure of the graph favors clustering state 4 together with state 3 since they belong to the same community. The higher-than-average reward of state 4 then drives up the average reward *θ* for that cluster, which in turn drives up average reward *μ* for the states that belong to it, and in particular for state 3. In contrast, state 7 often ends up in a separate cluster which is not influenced by the reward of state 3 and thus has an expected reward of 15, the average 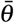 for the entire graph.

### Experiment seven: rewards and planning

While the results of experiment six validated the reward assumptions of our model, they could also be accounted for by alternative models, such as the successor representation (Stachenfeld et al., 2017; Dayan, 1993) or a Gaussian process with a diffusion kernel (Kondor and Lafferty, 2002; Wu et al., 2018). We therefore sought to test the unique prediction of our model that, in the absence of community or task structure, hierarchical planning will occur over clusters delineated by boundaries in the reward landscape of the environment.

#### Participants

We recruited 174 participants (68 female) from MTurk. Participants were paid $0.50 plus a bonus equal to the number of points earned on a randomly chosen trial in cents (up to $3.00). This encouraged participants to do their best on every trial. The experiment took 9 minutes on average.

#### Design

We asked participants to navigate a graph (“network of gold mines”) with the same structure as in experiments one and four (Figure 15A). Unlike the previous experiments, participants performed a mix of randomly shuffled free choice and forced choice trials. On free choice trials, participants started in a random node and could navigate to any node they chose. Once they reached their target node, they collected the reward (“grams of gold”) from that node. On forced choice trials, participants had a specified random target node and they could only collect reward from that node, similarly to the tasks in experiments one through five. Free choice trials encouraged participants to learn the reward distribution, while forced choice trials encouraged planning and prepared participants for the test trial, which was also a forced choice trial. Crucially, the rewards favored a clustering like the one in experiments one and four: states 1,2,3 always delivered the same reward, as did states 4,5,6 and states 7,8,9,10. Similarly to experiments one and four, participants were tested on a forced choice task from node 6 to node 1.

**Figure 15.**
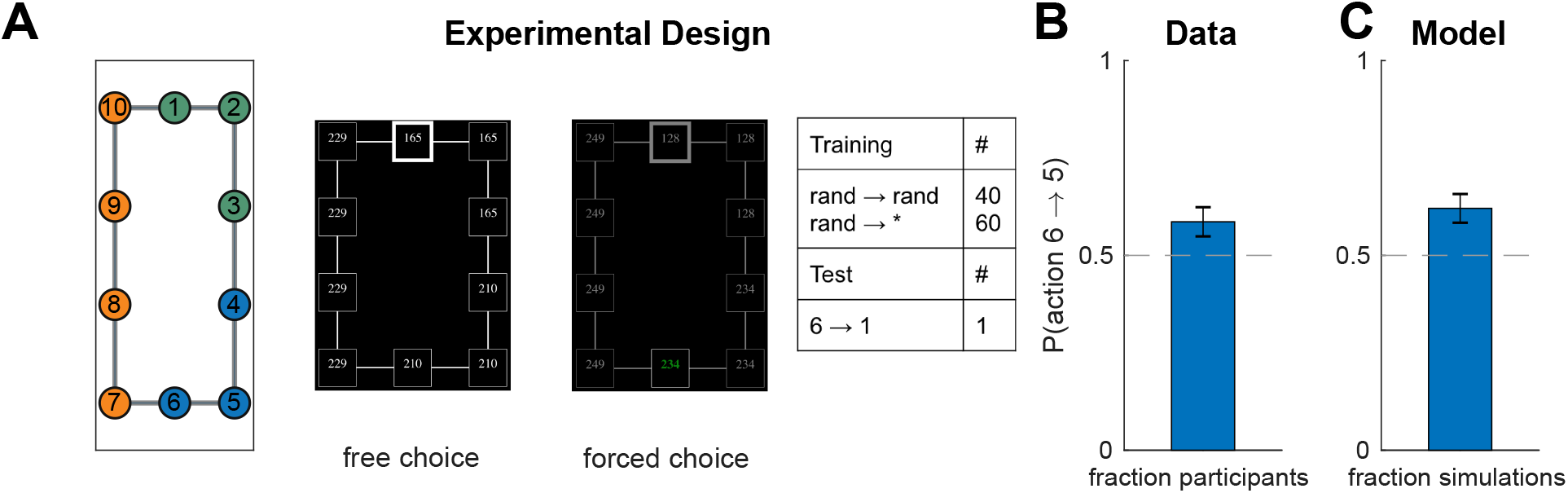
Rewards induce clusters that influence planning. A. (Left) Experiment seven employed the same graph as in experiments one and four, with the difference that clusters were induced via the reward rather than the task distribution. (Middle) screenshots from free choice and forced choice trials. (Right) training and test protocol. “Rand” indicates that a random state was chosen on each trial, while the asterisk indicates a free choice trial (i.e., the participant was free to choose any node). B. Results from experiment seven showing that participants were more likely to prefer the path with fewer reward cluster boundaries. Error bars are s.e.m. (174 participants). C. Results from simulations showing that the model exhibited the same preference. Notation as in B.

#### Procedure

Participants were told they would navigate a virtual network of gold mines. The graph layout was identical to experiment four (Figure 15A) and participants could see a bird’s-eye view of the entire graph, with each node representing a mine and each edge representing a tunnel between mines. Each mine delivered a certain number of points, which were displayed inside the node of the corresponding mine. Similarly to experiment four, participants could navigate between mines using the arrow keys and choose a mine using the space bar, after which they were informed how many points they earned from the chosen mine and immediately began the next trial. Mines always delivered deterministic points as indicated on each mine. Only points earned from the chosen mine counted.

On free choice trials, any mine could be chosen. On forced choice trials, only a single target mine was available and all other minds are grayed out (Figure 15A, right). Attempting to select a different mine resulted in a “incorrect” message flashing and did not change the current mine. There were 60 free and 40 forced choice training trials, starting at random states. The target state on the forced choice trials during training was also random. In order to prevent participants from developing a model-free bias towards one side of the graph based on reward magnitude (for example, because states there happened to be more rewarding), we randomly resampled the points of all states with probability 0.2 on every free choice trial. All reward values were sampled uniformly between 0 and 300.

#### Results and discussion

As in experiments one and four, participants showed a preference for the route with fewer cluster boundaries (Figure 15B; 102 out of 174 participants, *p* = 0.03, two-tailed binomial test). The model made same prediction (Figure 15C; 108 out of 174 participants, *p* = 0.001, two-tailed binomial test). However, this time the clusters were inferred solely based on rewards (Figure 15D), rather than topological structure or task distributions. This shows that the reward distribution in the environment can affect hierarchy discovery and consequently planning over that hierarchy, in accordance with our model predictions.

### Experiment eight: uncertainty and active learning

With the exception of experiment three, the results so far could also be accommodated by a non-Bayesian hierarchy discovery account that simply searches for the “best” hierarchy as defined by some utility or score function (Fernández and González, 2013). Indeed, our inference algorithm could be viewed as a form of stochastic search over the space of possible hierarchies, with the (unnormalized) posterior simply serving the role of a score function for comparing candidate hierarchies. Our hierarchical planner also relies on a single point estimate of the hierarchy, which raises the question of whether a Bayesian account is warranted. While the results of experiment three show that the uncertainty in the posterior can affect choices across participants (since the population can be seen as approximating the posterior, with one sample hierarchy per participant), they do not address the question of whether uncertainty is represented at the level of the individual participant.

In this experiment, we sought to explicitly validate the probabilistic aspects of our model by showing that people can make choices based on the uncertainty of the posterior distribution over hierarchies *P*(*H*|*G*). Previous work has demonstrated that both humans and animals have access to their uncertainty (Smith et al., 2003), which they can report explicitly or leverage to direct their exploratory choices towards informative options (Gershman, 2018). Most related to our study is work on active learning (Bramley et al., 2015; 2017) showing that people choose to learn in a way that maximally reduces their uncertainty about the hidden structure of the environment. Applied to our model, this predicts that people would choose to learn about aspects of the graph that maximally reduce their uncertainty over the hierarchy, which we operationalize as the entropy of the posterior (see Appendix).

#### Participants

We recruited 40 participants from the Harvard community. The experiment took around 5 minutes and participants were not paid for their participation.

#### Design

We presented 40 participants with 7 graphs (“subway networks”; Figure 16A). In each graph, all but two or three of the edges were observable (those edge were described as “tracks possibly under construction”). We then asked participants to indicate which of the currently unobservable edges they would like to observe (i.e., to learn whether the edge is present or not). Participants could only pick a single edge for each graph.

**Figure 16.**
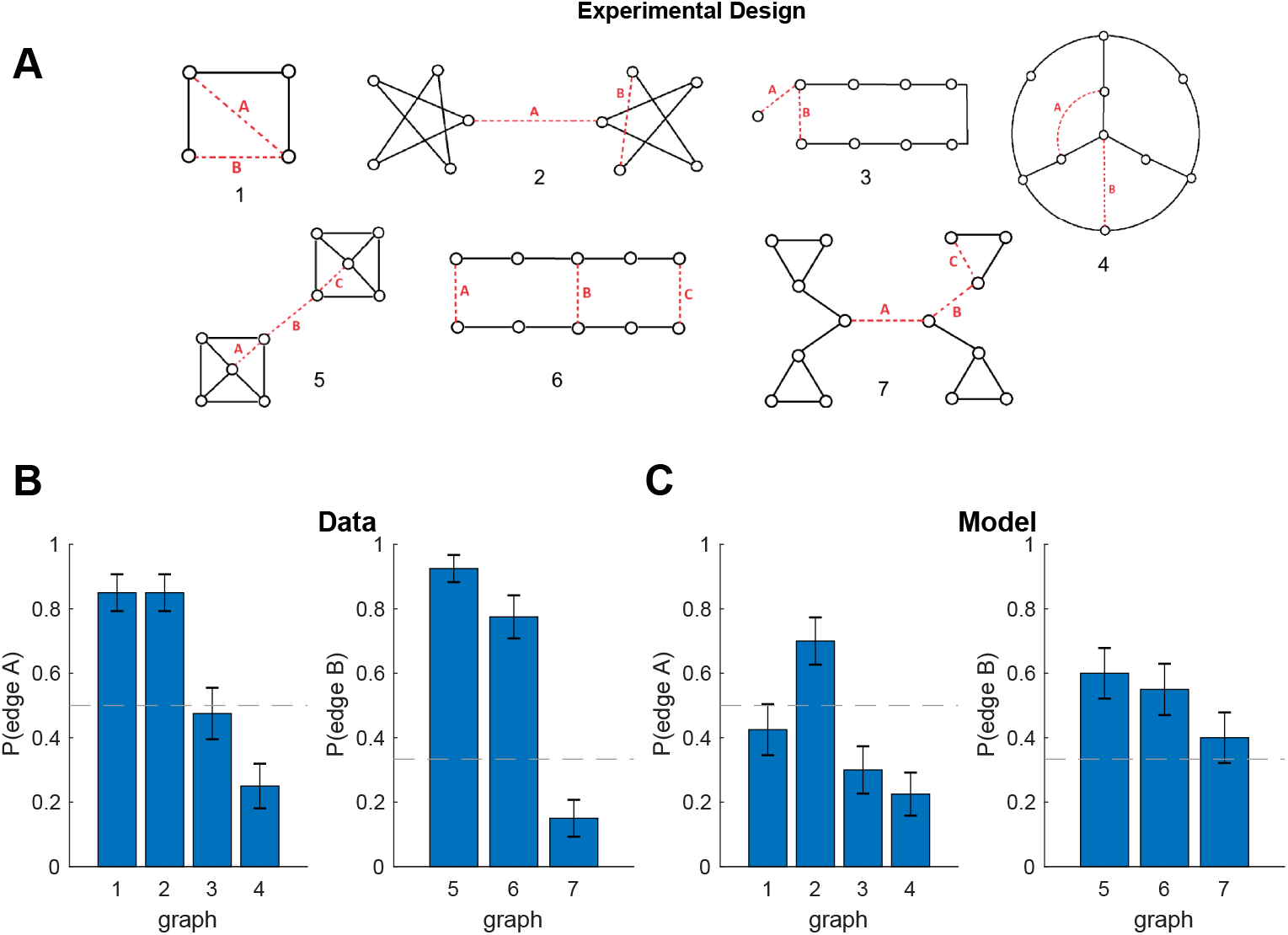
Uncertainty about hierarchy guides active learning. A. The seven graphs shown to participants in experiment eight. Dashed lines indicate edges whose status was unknown. Numbers are graph identifiers. B. Results from experiment eight showing what fraction of participants chose to learn about edge A in graphs 1-4 (left) and what fraction chose to learn about edge B in graphs 5-7 (right). Dashed line is chance. Error bars are s.e.m. (40 participants). C. Results from simulations showing that the model exhibited similar preferences. Notation as in B.

#### Procedure

Each participant was given a sheet of paper with instructions and the graphs in Figure 16A. The graphs were presented in a different order. The instructions were as follows:

*Imagine that you are a tourist visiting a new city and want to use the subway system to get around. Unfortunately, the subway map you have is out of date, and several tracks on it are marked as “Under Repair”. As it might help get you around faster, you decide to ask a passer-by if any of these tracks have finished repair work. Each of the images below represent a subway network, with the circles as stations and lines as tracks. For each subway system, there are 2 or 3 tracks in red, representing tracks that are possibly under construction. Circle ONE track whose repair status you want to find out about the most. There is no need to think too hard about each map!*

#### Results and discussion

If participants only care about the flat graph G, then they should be indifferent between the available choices since they all yield exactly one bit of information. Furthermore, if participants maintain a single point estimate of the hierarchy without keeping track of uncertainty, they would have no basis to estimate how informative each choice is for the purposes of hierarchy discovery. In contrast, we found that participants showed a strong preference in 5 of the 7 graphs (Figure 16B; *p* < 0.02 for graphs 1,2,4,5,6; binomial tests, Bonferroni corrected for 7 comparisons).

The simulated choices of our model were in agreement with the empirical results (*r* = 0.76, *p* = 0.05, Pearson correlation). This suggests that people explore their environment in a way that facilitates hierarchy discovery, in accordance with our Bayesian account. The main discrepancy between our model predictions and the participants’ choices was for graphs 1 and 7. This is likely due to other factors also playing a role in human choices, such as graph geometry or connectivity. In addition, since the model parameters were tuned based on simulations of previous studies that did not assess the effects of uncertainty, it could be that the model weighs various aspects of the graph differently from participants when computing the posterior, which would result in different uncertainty estimates. We leave the fine-tuning of model parameters as the subject of future work.

### Neural implementation

Finally, we speculate about how the computational processes proposed here might be implemented in neural circuits, and which brain regions might perform the computations. We also simulate within-trial and across-trial neural signals that could be used to identify the key brain areas involved in hierarchy discovery and hierarchical planning.

First, consider the flat graph *G* only. A straightforward way to encode *G* in a neural circuit would be to have a single unit (a neuron, such as a place cell, or an ensemble of neurons) represent each node *u* ∈ *V* and excitatory synapses between pairs of units (*u, v*) represent the edges *E* (Figure 17A, bottom). The graph structure could be learned via local Hebbian plasticity: when two units are activated right after each other (for example, during a transition between the corresponding states), the synapse between them is potentiated. In order to perform (flat) BFS to find the shortest path from node *s* to node *g*, an external input can successively probe each neighbor of *s* by transiently activating the corresponding unit, triggering a “forward sweep” of activation that propagates through the circuit until it reaches and activates the unit of the goal state *g*. The neighbor of *s* that activates *g* in the least amount of time is the next node along the shortest path to *g*, so the agent can then physically transition to that state and repeat the process again and again, until finally reaching the goal state g. Assuming some form of short-term synaptic depression or ion channel inactivation that prevents the sweep from going backwards (Dobrunz et al., 1997), this implements precisely BFS. Similar schemes have been proposed to support forward trajectory planning in the hippocampus (Erdem and Hasselmo, 2012; Gönner et al., 2017).

**Figure 17.**
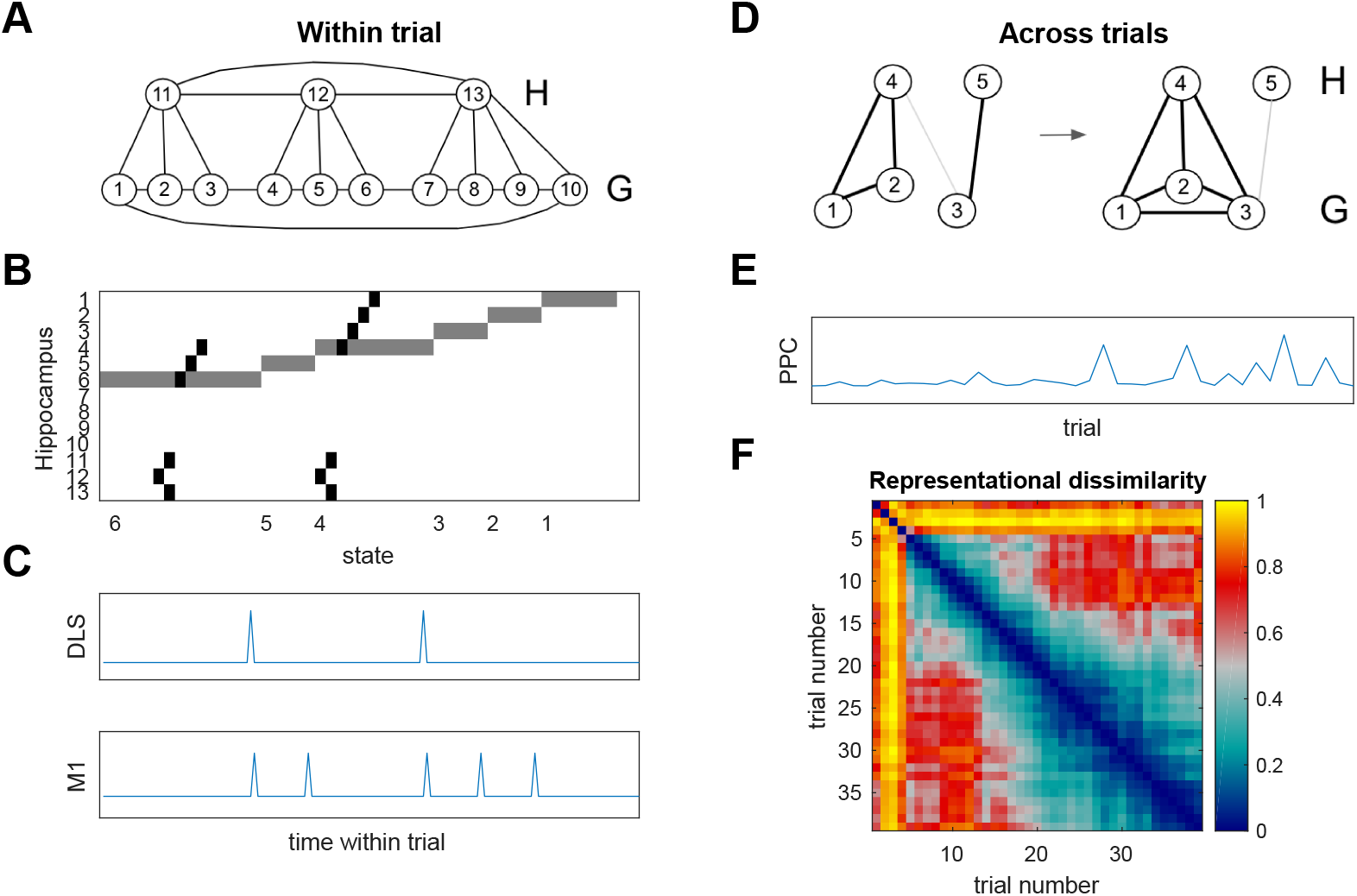
Hierarchy discovery and hierarchical planning in the brain. A. Example neural circuit encoding the low-level graph *G* (bottom), the high-level graph *H* (top), and the cluster assignments *c* from experiment one (Figure 9A). Circles denote units representing graph nodes. Lines denote bidirectional excitatory synapses representing edges and cluster assignments. Number are unit identifiers. B. Example (idealized) circuit activity during the test trial 6 → 1. Each row represents the activity of the corresponding unit over the course of the trial. States along the X-axis denote the current state following a transition. Gray denotes intermediate levels of activation representing the current state of the agent, akin to hippocampal place cell activity. Black denotes high levels of activation during planning, akin to hippocampal preplay. C. (Top) Example (idealized) “start” activity at the initiation of each action chunk in dorsolateral striatum (DLS), with action chunks assumed to fall within the boundaries of the state chunks (clusters) in A. (Bottom) For comparison, primary motor cortex (M1) activity at key presses corresponding to transitions. Time course corresponds to B. D. Example neural circuit illustrating hierarchy discovery via local Hebbian plasticity. (Left) Low-level graph with a single edge (1,2) has nodes 1 and 2 assigned to cluster 4 and node 3 is assigned to cluster 5. (Right) Observing edges (1,3) and (2,3) causes transient activation of nodes 1,2,3 and cluster 4, strengthening the connection between node 3 and cluster 4 and hence reassigning node 3 to cluster 4. E. Simulated Bayesian update of the (approximate) posterior *P*(*H*|*D*) over the course of learning the graph from simulation four (Figure 7A), which could take place in posterior parietal cortex (PPC). F. Representational dissimilarity matrix showing the difference in the (approximate) posterior *P*(*H*|*D*) between pairs of trials during the same simulation as in E.

The hierarchical graph *H* could then be incorporated into the circuit by designating its own set of units and synapses, corresponding to the nodes *V*′ and edges *E*′, respectively. The cluster assignments *c* could also be implemented as synapses between the units of *G* and the units of *H*. In an elegant way, the inferred hierarchy would be isomorphic to the neural circuit that represents it (Figure 17A). This could straightforwardly extend to deeper hierarchies and is consistent with the presence of place cells and grid cells with different receptive field sizes in the hippocampus and entorhinal cortex (Jung et al., 1994; Kjelstrup et al., 2008). HBFS can be implemented in the exact same way as BFS, with the difference that now the forward sweep can take “shortcuts” through the higher levels of the hierarchy, thus significantly reducing the time it would take to activate the goal unit *g*. Note that this performs almost the exact same computation as HBFS, with the small difference that transitions “up” the hierarchy (i.e., computing *c_u_* and *c_v_* on line 1 in Algorithm 1) also count as transitions along the path.

The hierarchy could be learned similarly to *G*, with local Hebbian plasticity strengthening the synapses between units of *H* (for example, during boundary transitions) as well as the synapses representing the cluster assignments between *G* and *H*. To illustrate this, consider the example in Figure 17D (left), with units 1,2,3 representing nodes in *G* and units 4,5 representing nodes in *H*. On the left, there is only one edge between nodes 1 and 2 and hence these two nodes are clustered together (*c*_1_ = *c*_2_ = 4), separately from node 3 (*c*_3_ = 5). However, once the edges (1,3) and (2,3) are observed (Figure 17D, right), this would transiently activate units 1,2,3 together, which (because of nodes 1 and 2) would transiently activate unit 4, leading to potentiation of the synapse between 3 and 4. Assuming some form of local homeostatic plasticity that constrains the total synaptic weight of each unit, this would weaken the synapse between 3 and 5, effectively reassigning 3 to the same cluster as 1 and 2 (*c*_1_ = *c*_2_ = *c*_3_ = 4).

Note that this implements a kind of “soft” hierarchy, with the same node potentially being strongly associated with one cluster and weakly associated with other clusters. This could be one way to take into account a probability distribution over hierarchies rather than a single point estimate. Indeed, allowing all synaptic weights to have continuous values rather than forcing them to be binary can keep track of probability distributions over the edges *E* and *E*′ as well. In fact, this could also allow the neural circuit to take into account stochastic transitions: transitions that have low probability will simply have the corresponding synapses potentiated less often, resulting in weaker weights. This addresses the limitation of our graph-theoretic approach to only support deterministic transitions, thus extending the framework to support regular MDPs. The continuous range of the synaptic weights would naturally be taken into account by our “neural” HBFS algorithm: the forward sweep will simply be less likely to propagate through weaker synapses. In effect, this will perform a kind of simultaneous, parallel sampling of an entire set of possible trajectories in a way that is drastically more efficient than sampling trajectories one by one (linear versus exponential time). Investigating the theoretical properties of such a mechanism could be the subject of future work.

A neural circuit with the above-mentioned properties could naturally be implemented in the hippocampus and the surrounding cortex. Hippocampus has long been known to encode locations in physical space (O’Keefe, 1976) and has been hypothesized to encode a cognitive map that applies across various non-spatial domains (O’Keefe and Nadel, 1978). Recent studies have shown that this is indeed the case, with encoding of non-spatial task-relevant variables such as sound frequency in rodents (Aronov et al., 2017) and even abstract conceptual domains in humans (Constantinescu et al., 2016). The units we hypothesize could thus be implemented in the hippocampus-entorhinal circuit, with HBFS taking the form of hippocampal preplay (Figure 17B) which is known to occur at decision points and is predictive of future behavior (Dragoi and Tonegawa, 2011).

While our model only discovers state chunks, action chunking could straightforwardly be incorporated by caching (or memoizing) the output of BFS and/or HBFS for regularly occurring subgoals (see Future directions). The acquisition of action chunks after extensive training in animals is associated with the emergence of characteristic start/stop signals in basal ganglia circuits (Graybiel, 1998; Barnes et al., 2005; Jin and Costa, 2010; Jin et al., 2014; Geddes et al., 2018). Our model makes the distinct prediction that action chunks and the corresponding start/stop signals will fall within state chunk boundaries (Figure 17C), rather than be dictated purely by the temporal statistics of action sequences. This prediction could be validated empirically by subjecting animals to similar training and test protocols as our participants while measuring neural activity in dorsolateral striatum.

If the probability distribution over hierarchies is implicitly encoded in synaptic weights, as proposed above, then it would be difficult to read it out directly from neural activity. Alternatively, the distribution could be encoded in neural activity patterns, for example using probabilistic population codes or neuronal sampling (Pouget et al., 2013). This would be consistent with our previous work (Tomov et al., 2018) which found a neural signature of the Bayesian update of the posterior over hidden structures in a frontroparietal network of brain regions, as well as representations of the full posterior in several brain areas. Our model falls within the framework of structure learning and is thus likely to recruit the same underlying neural mechanisms. This prediction can be tested by generating regressors that track Bayesian updates of the posterior *P*(*H*|*D*) during learning (Figure 17E) and using them to identify neurons or brain areas that might implement the actual hierarchy discovery process. In addition, representational similarity analysis (Kriegeskorte et al., 2008) could be used to identify areas that maintain the (approximate) posterior *P*(*H*|*D*) (Figure 17F).

## General Discussion

In this study, we proposed a Bayesian model for discovering state hierarchies based on a generative model of the topological structure, rewards, and tasks in the environment. Building on the Bayesian brain hypothesis (Dayan et al., 1995; Chater et al., 2006; Clark, 2013) and on principles developed in structure learning (Gershman et al., 2015) and robotics (Fernández and González, 2013), it postulates that the brain learns useful hierarchical representations by inverting this generative model to infer the hidden hierarchical structure of the world. These representations can be subsequently used to plan efficiently and flexibly in the face of changing task demands. The model accounts for a number of phenomena previously reported in the literature and makes new predictions which we verified empirically.

The model goes beyond previous accounts of clustering states in the environment both in the scope of findings it can explain and in the predictions it makes. Schapiro et al. (2013) proposed a recurrent neural network that learns community structure based on the temporal statistics of the environment. Their model and other event segmentation models (Zacks and Swallow, 2007) can learn to predict stimuli that tend to co-occur, such as states in the same community, and can detect boundaries when those predictions are violated, for example after a transition to a new community. While this could explain the effects of simulations one through five and possibly experiments one and two, it will be challenged by experiments four, five, six and eight, in which the environment is fully visible in all stimuli co-occur together. More importantly, these models make no predictions regarding action selection, and it is not clear how the learned representations can be used for planning. Solway et al. (2014a) offer a formal definition of an optimal hierarchy; however, as the authors themselves point out, their analysis is not a plausible account of how hierarchy is learned, because it assumes the agent already knows the optimal solution to all possible tasks in the environment, which defeats the purpose of learning a hierarchy in the first place. A detailed comparison between our model and alternative candidate models that make similar claims is shown in Table 2.

**Table 2.**
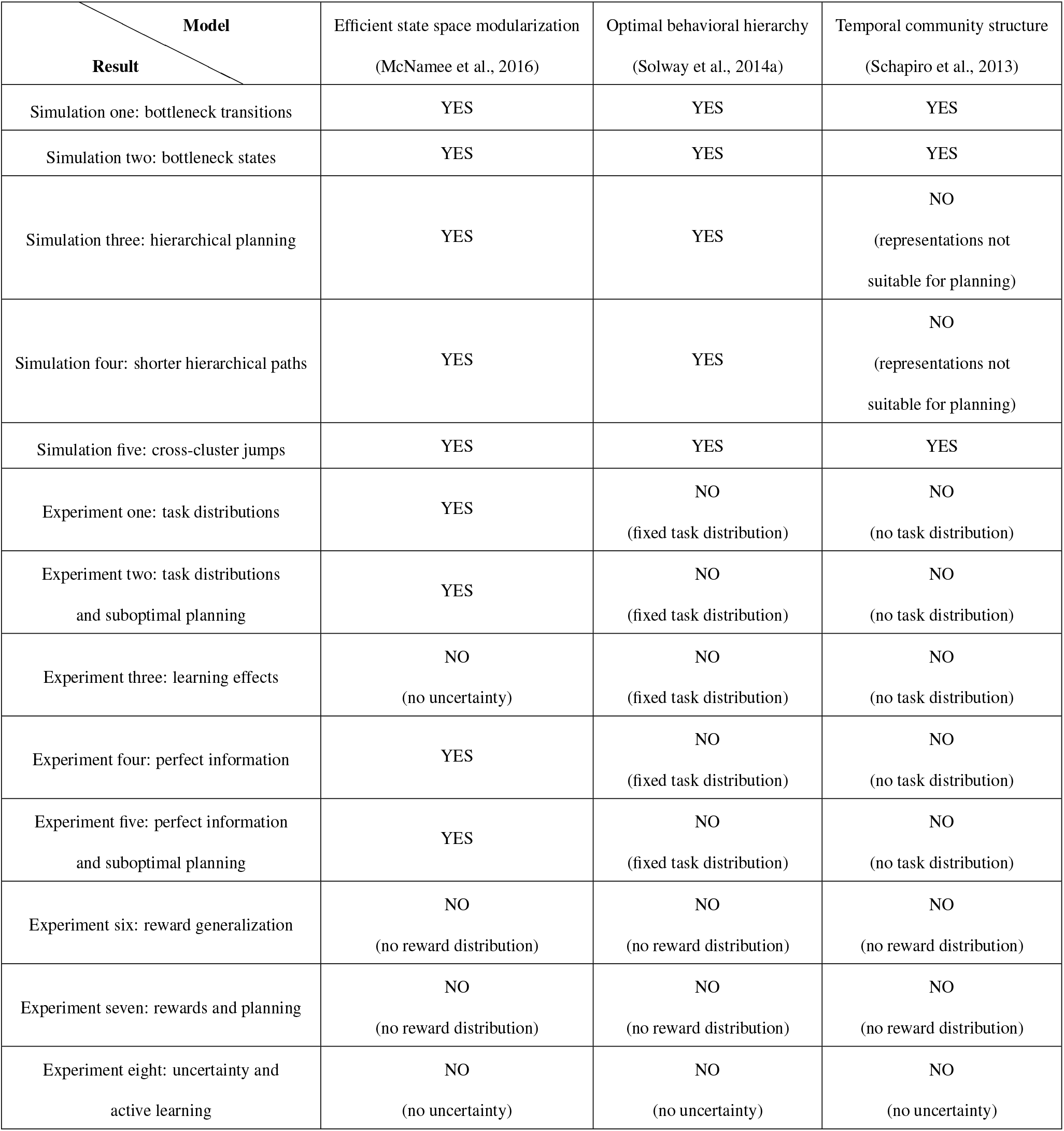
Model comparison. Summary of which results could potentially be accounted for by alternative models and which results rule out certain models.

The problem of planning has been studied extensively both for biological as well as artificial agents. Correspondingly, our work resonates with several important strands of research, which we discuss in turn below.

### Cognitive architectures

The earliest attempts to develop a formal theory of planning date back to the work of Newell and Simon (Newell et al., 1958; 1972) who laid the foundational concepts of planning and problem-solving both in human and in artificial intelligence research. Simon (1978) framed problem-solving as an interaction between the participant and the environment, and planning as a search through a state space that represents the structure of the problem, with operators (or actions) performing transitions between states. In parallel, the seminal work of Miller et al. (1960) highlighted the hierarchical organization of human action plans, which they linked to people’s highly structured representations of the world. Miller et al. (1960) also proposed the existence of a fast-access, limited-capacity working memory that loads information from a large-capacity “dead storage” system, concepts later formalized as the short-term and long-term memory stores in Newell et al. (1972)’s production systems. These earlier attempts led to the development of contemporary cognitive architectures such as Soar (Newell, 1992; Laird, 2012) and ACT-R (Anderson, 1993; Anderson et al., 2004), which aim to capture all aspects of human cognition. Both of these systems can perform hierarchical problem-solving in complex domains based on subgoals, however the subgoals have to be supplied manually. Chunking (referred to as compilation in ACT-R) is implemented by caching or memoizing the solutions to these subgoals (Laird et al., 1986).

Our approach builds on concepts developed in this tradition. In our model, hierarchical behavior directly arises from the hierarchical representation of the environment. The two memory systems we assume also mirror the short-term/long-term memory stores in these earlier cognitive accounts. Additionally, the form of action chunking we propose as a future addition to the model (see Future directions) is similar in spirit to the chunking mechanisms in Soar and ACT-R. In principle, our hierarchy discovery method could be integrated with these production systems to allow them to decompose the problem space and identify subgoals automatically.

### Model-based and model-free reinforcement learning

In psychology, planning and goal-directed behavior, which selects actions based on their outcomes, is often contrasted with habitual behavior, which selects actions automatically (Dolan and Dayan, 2013). The dichotomy between these two kinds of action selection strategies – fast, reflexive, habitual, unconscious responding (also referred to as System 1) on one hand, and slow, reflective, goal-directed, conscious deliberation (also referred to a System 2) on the other – has permeated the field for over a century (Thorndike, 1911; Tolman, 1948), has been reincarnated in many forms (Dickinson et al., 1983; Kahneman and Egan, 2011; Stanovich and West, 2000), and is realized in distinct neural circuits (Yin et al., 2004; 2005; Balleine and Dickinson, 1998; Balleine, 2005). In RL, habitual and goal-directed behaviors have been associated with *model-free* and *model-based* algorithms, respectively (Doya et al., 2002; Daw et al., 2005).

In model-free algorithms, the agent learns a value function for each state and/or each action by trial-and-error, and then chooses actions with respect to the learned values. In model-based RL, the agent learns the reward and transition structure of the environment and combines that information to find actions leading to reward. Model-free RL is associated with fast, reflexive responding because it myopically considers the value of the current state/action only, while model-based RL is associated with slow, reflective responding because it considers the transitions and rewards of multiple states/actions ahead. Despite the superficial similarity of these two systems and the two computational components we propose – a fast online planner, and a slow offline representation learner – there is a profound difference. Both model-free and model-based RL (as well as other versions of habitual and goal-directed strategies) perform online action selection, with agents having to rely on one or the other to solve a particular task. In contrast, our planner makes decisions about which actions to select in a particular task, while the hierarchy discovery process works offline, outside the context of a particular task. In a way, our planner is more like model-based RL in that it performs deliberate, goal-directed computations based on its internal model of the world (the “model” in model-based RL). The purpose of the hierarchy discovery process is to learn such an internal representation that can be used by a planner with limited computational resources. Indeed, the cognitive limitations inherent in this type of reflective decision-making (Kahneman and Egan, 2011) are the foundational assumptions behind our approach.

Notice that we only invoke a model-based planner in order to motivate hierarchy discovery and to link the hierarchy to decision-making, without committing to the particular HBFS algorithm. While hierarchy discovery can be justified in this way without invoking a model-free system, we certainly do not exclude the possibility of such a system existing and operating in parallel with the planner. A model-free component could easily be incorporated into our framework, for example by having a parallel model-free system which learns the values of states and actions. During decision making, the agent can use a meta-cognitive process to arbitrate between the planner and the model-free system (Kool et al., 2017). Like the planner, the model-free system could also operate at different levels of abstraction, learning values for the clusters in addition to the low-level states. This could account for effects of reflexive responding that we explicitly controlled for in our experiments.

Another distinguishing feature of our approach is that, by leveraging the deterministic nature of the transition structure, our planner can rely on simple shortest path algorithms to find solutions to tasks. In contrast, traditional approaches to model-based RL involve computationally costly operations such as value iteration or sampling of candidate trajectories, both of which scale poorly with the size of the state space. Even though the combinatorial explosion of sampling could be managed by heuristics such as pruning of decision trees (Huys et al., 2012; 2015), such approaches are unnecessary in the deterministic domains often considered in planning problems.

It is worth highlighting that our results cannot be explained by standard model-free or model-based RL algorithms. In experiments one, two, four, and five, model-free RL would have learned high values for states 3, 6, and 7, and hence would prefer the transition 6 → 7 on the test trial. In experiments one and four, model-based RL would be indifferent between the two possible trajectories on the test trial due to the symmetry of the graph. In experiments two and five, model-based RL should actually favor the shorter path via 6 → 7. All of these predictions go against our participants’ tendency to pick the transition 6 → 5.

The two systems in our proposal might also bear superficial resemblance to the Dyna architecture in RL (Sutton, 1991) in which a slow, offline model-based simulator trains a fast, online model-free system to respond adaptively to situations it has never experienced before. This process is reminiscent of hippocampal replay (Shohamy and Daw, 2015) and is particularly useful when the environment changes too often for trial-and-error learning to be effective. In contrast, we propose an offline inference process for learning representations (the “model” in model-based RL) that a model-based system can use to plan in previously unseen tasks. If our proposal is extended with a separate model-free component as suggested above, it could be integrated with Dyna, which would in turn use the model-based system to train the model-free system.

### Hierarchical reinforcement learning

A long-standing challenge for traditional RL has been the combinatorial explosion that occurs when planning and learning take place over long time horizons. This challenge been addressed by hierarchical RL (HRL), which breaks down the problem into sub-problems at multiple levels of abstraction. One influential approach to HRL extends the agent’s action repertoire to include *options* (Sutton et al., 1999) which consist of sequences of actions (the option policy) that accomplish certain subgoals (for example, exiting a room). When an option is selected, the corresponding action sequence is executed as a single behavioral unit. Options are also referred to as skills, subroutines, partial policies, macro-actions, or policy chunks, while the original actions are sometimes referred to as primitive actions.

As with regular or “flat” RL, HRL also comes in two distinct flavors (Botvinick et al., 2009): model-free HRL and model-based HRL. Like model-free RL, model-free HRL (Dezfouli and Balleine, 2012; Holroyd and McClure, 2015; Rasmussen et al., 2017) learns a value function; however, in this case the value function is additionally defined for options. Error-driven learning occurs both on the high level by learning which options lead to rewarding outcomes, as well as on the lower level by learning the best option policy for each option. Importantly, there is no notion of a transition function which could be used for planning, and hence model-free HRL could not exhibit the behaviors predicted by our model. Although a model-free component could be incorporated into our framework as discussed above, it would not account for the results of experiments one through five. The only options that could conceivably have been learned by model-free HRL are the ones corresponding to the training tasks (for example, 1 → 3, 4 → 6, and 10 → 7 for experiment one). This set of options could not explain the results of the test trial which requires going in the opposite direction.

In model-based HRL (Botvinick and Weinstein, 2014), as in model-based RL, the agent separately learns a transition function and a reward function. Additionally, the agent is furnished with an *option model* that specifies the initiation states (for example, locations within a room), termination states (subgoals; for example, doors), average duration, and average reward of each option. The agent can thus plan over options rather than primitive actions, essentially performing mental “jumps” in the state space, allowing it to first form a high-level plan between subgoals reachable via options and then refining that plan on the lower level by simply following the corresponding option policies. This form of *saltatory* model-based HRL is conceptually identical to our proposal. While our work builds on concepts developed in parallel to HRL in the field of robot navigation and planning (Fernández and González, 2013), our model can be cast in HRL terms by considering each task as equivalent to placing a positive reward in the goal state and a small negative reward in all other states, thus encouraging the agent to find the shortest path to the goal state. Edges in the high-level graph *H* can be seen as options, with subgoals specified by the endpoints of bridges and option policies specifying how to reach the subgoals within a cluster.

Our work introduces two critical improvements to model-based HRL. First, as in model-based RL, model-based HRL assumes planning occurs by sampling trajectories through the state space, which in our proposal is performed by the deterministic and much more efficient HBFS algorithm. Second, as Botvinick and Weinstein (2014) point out, a critical open question is how useful options are discovered in the first place. The approach they propose is based on the successor representation (Dayan, 1993) which partitions the state space along topological bottlenecks. However, this would not predict the results of experiments one through five and experiment seven, in which clustering occurs based on tasks and rewards only.

When framed within HRL, our approach can be viewed as a solution to the option discovery problem which has plagued the field of HRL since its inception, as the original formulation never specified how useful options are learned in the first place. Discovering useful subgoals and, correspondingly, useful options is critical, since an inadequate set of options can lead to dramatically worse performance compared to regular RL (Botvinick et al., 2009). This has led to the proliferation of a rich literature on option discovery, to which we turn next.

### Option discovery

While earlier work on HRL assumed the options are supplied manually (Sutton et al., 1999), a growing number of HRL studies have focused on the problem of discovering useful options. A detailed review of the option discovery literature is beyond the scope of this discussion, but we highlight some of the main approaches. Most option discovery methods fall in one of two broad categories: *state abstraction* and *temporal abstraction* methods (McGovern, 2002).

State abstraction methods first decompose the state space by identifying clusters or subgoals, and subsequently identify options based on that decomposition (Dayan and Hinton, 1993; Vezhnevets et al., 2017). The state space could be partitioned into clusters based on the value function (Dietterich, 2000), the state features (Hengst, 2002), or the transition function (Machado et al., 2017; Ravindran and Barto, 2002). Other approaches designate certain states as subgoals and learn options that lead to those subgoals, which could be defined by salient events (Chentanez et al., 2005), object-object interactions (Kulkarni et al., 2016), frequently visited states (Stolle and Precup, 2002; McGovern and Barto, 2001; Digney, 1996), or large changes in the reinforcement gradient (Digney, 1996). Other subgoal discovery methods rely on graph theoretic notions to identify bottlenecks (such as “doors” between rooms; Simşek and Barto, 2008), the boundaries of strongly connected regions of the state space (such as the “walls” of rooms; Menache et al., 2002), or clusters of states (such as the rooms themselves; Mannor et al., 2004; Şimşek et al., 2005) as subgoals.

Temporal abstraction (or policy abstraction) methods directly learn the option policies, without resorting to state abstraction as an intermediate step. Some of these approaches identify frequently used action sequences from successful trajectories (McGovern, 2002; Girgin et al., 2006; Vezhnevets et al., 2016). Other approaches posit a generative model for policies that favors temporal abstraction, and then perform probabilistic inference to find the optimal policy (Wingate et al., 2013; Daniel et al., 2016).

Viewed in HRL terms, our model falls into the state abstraction category, since it first partitions the state space into clusters which in turn define subgoals and constrain behavior. However, our model goes beyond these previous attempts: it unifies multiple ideas from these different approaches under a single Bayesian framework that allows it to account for all the behavioral phenomena in our study. To the best of our knowledge, no single option discovery method based on state abstraction would capture all of them. Option discovery methods based on temporal abstraction would also fail to account for the results of experiments one through five, since participants were never trained to navigate in the directions tested in the test trial.

### Information-theoretic approaches

Closely related to our study is work by McNamee et al. (2016) proposing an alternative approach to hierarchically decomposing the environment for planning under working memory limitations. Similarly to our proposal, they divide the state space into clusters (or modules) and assume planning first occurs at a high-level (across modules), and is subsequently refined at a lower level (within modules). They define an optimal modularization of the state space as the one which minimizes the expected information-theoretic description length of planning trajectories. Intuitively, this means that the average hierarchical plan in the modularized state space is as simple as possible. Unlike the analysis of Solway et al. (2014a), this method does not require knowing the optimal behaviors in advance and can also accommodate different task distributions. This implies that it could account for effects based on graph topology and task distribution (see Table 2). Unlike our model, it does not account for effects of the reward distribution and uncertainty. By framing the process of hierarchy discovery in terms of Bayesian inference, our model can be used to make predictions about how beliefs evolve during learning (see experiment three and Future directions), and how plans and choices will change correspondingly. Performing Bayesian inference incrementally in this way can also be used to investigate the neural correlates of hierarchy discovery and to understand the underlying neural computations (Figure 17E,F). Indeed, our model does not appeal to a strict definition of optimality (in the sense of producing a hierarchy that is provably optimal), and hence the two approaches can be seen as complementary, with our model explaining how hierarchy is discovered and the analysis of McNamee et al. (2016) validating the hierarchies learned by our model and by people.

Another information-theoretic method for clustering state spaces was proposed by Maisto et al. (2016). Their approach relies on an extension of the CRP that allows clustering based on similarity between states, which can be defined via a prespecified kernel function. Using different kernels, they demonstrate clustering based on bottlenecks, goal states, paths, or other aspects of the graph structure. Their approach relies on algorithmic probability theory to define the kernels (see also Donnarumma et al., 2016). This involves precomputing all possible paths between each pair of states, which renders planning unnecessary. Nonetheless, this approach could provide a useful tool to analyze the optimality of the hierarchies inferred by our model.

### Structure learning and other notions of hierarchy

Frank and Badre (2011) proposed an alternative notion of hierarchy in terms of action rules at different levels of abstraction (see also Collins and Frank, 2013; 2016). In their framework, low-level rules map stimuli to responses, whereas high-level rules dictate which stimulus dimensions are relevant for the low-level rules. For example, a low-level rule might say “if the traffic light is red, don’t walk”, while a high-level rule might say “when crossing the street, pay attention to the color of the traffic light”. This implements a form of *state aggregation*, which in RL refers to the grouping together of different states and then treating them as a single state, for example by assigning the same values and actions to all states in the group (Moore, 1991; Singh et al., 1995). Note that this is different from state clustering in our model, which still treats each state within the cluster as distinguishable from the rest. By determining which stimulus dimensions are relevant for responding, the high-level rules implicitly render all states with the same value for the particular stimulus dimension as indistinguishable for the purposes of responding according to low-level rules (for example, “a red light on the sunny day” versus “a red light on a rainy day” both elicit the same response). This drastically reduces the total number of stimulus-response mappings (low-level rules) that need to be learned and allows generalization to previously unseen stimuli.

The connectionist model proposed by Frank and Badre (2011) implements error-driven learning at these different levels of abstraction in parallel loops that map onto the corticostriatal hierarchy, with more anterior regions representing rules at increasing levels of abstraction. Despite its ability to account for a range of behavioral and neural data, this approach is fundamentally restricted to learning stimulus-response mappings only, and as such falls into the category of model-free RL since it has no notion of a transition structure over which to plan. As discussed previously, model-free RL – even with state aggregation – cannot account for the results of experiments one through five since the predominant response (6 → 5) was never reinforced, nor would it support the kind of goal-directed planning presented here. In fact, state aggregation would likely not be possible in these experiments, since there is only a single stimulus dimension (the name of the current station). In their model, the term hierarchy is used to denote a hierarchy of rules that amounts to a compressed mapping from one stimulus to one action. This is fundamentally different from our notion of a hierarchy, in which a hierarchy of states supports the flexible generation of multistep action plans that achieve distant goals. It also differs from the traditional notion of HRL, which refers to a temporal hierarchy, with options consisting of sequences of primitive actions. While in theory the notion of a high-level rule could be extended to include options, the model they propose can only learn to ignore stimulus dimensions and as such can only compress stimulus-response mappings, whereas to the contrary, options require an expansion of the stimulus-response space.

### Partial observability

Closely related to state discovery models is work in RL on partially observable environments (Kaelbling et al., 1998). In this scenario, the agent never directly observes its current state but must instead infer it from observations as it interacts with the environment. Formally, the environment is represented by a partially observable Markov decision process (POMDP) in which states, actions and rewards are represented similarly to MDPs, with the key difference that states additionally generate observations. The agent then uses these observations to infer a probability distribution over states – the *belief state* – which it uses for decision making. Building on RL and Bayesian principles, POMDPs provide a normative way to maximize reward under uncertainty. Correspondingly, they have been used to account for a wide range of behavioral and neural results in the animal learning and decision making literature (Dayan and Daw, 2008; Rao, 2010). Recently, neurophysiological evidence from rodents (Starkweather et al., 2017; 2018; Babayan et al., 2018) has shown that midbrain dopaminergic firing is consistent with a RL signal computed over such a belief state, thus grounding the POMDP framework in the well-established brain circuits for reward-based learning.

Our model can be seen as an extension of the POMDP framework, with clusters (high-level states in *H*) acting as hidden states and low-level states in *G* acting as observations. However, our model differs from the standard POMDP definition in two ways. First, as latent cause models, POMDPs assume observations are independent given the state, whereas our model relies on the relations between states (*E*) in order to infer the clusters. Second, unlike latent cause models, POMDPs usually assume a prespecified state space, whereas our model allows for a theoretically unbounded number of clusters, recruiting more clusters as dictated by the data. The second property of our model makes it similar to an infinite POMDP (Doshi-Velez, 2009), which dynamically expands the state space as more observations are acquired. The first way that our model differs from POMDPs suggests that the analogy between observations and low-level states might be inappropriate, and that our model can be better thought of as a particular kind of infinite hierarchical MDP, in which states are fully observable but there is additional hidden structure which is not observable. Viewed in this way, our model does not support partial observability, a limitation which could be remedied by making the low-level states unobservable and having them generate observations, which would drive inferences about the states, which would in turn drive inferences about the clusters. While this would complicate the inference process, it would bring our model more closely in line with the POMDP framework, making it more applicable in a world in which agents only receive partial information about their state in the environment.

### Action chunking and motor sequences

Our work is closely related to the notion of hierarchical control and motor sequencing (Rosenbaum et al., 1983; 1984; Koch and Hoffmann, 2000), which studies the behavioral effects predicted by hierarchical action plans. Our work speaks directly to that literature by proposing one particular way in which the representations that support such hierarchical planning might be learned. Indeed, our hierarchical planner is reminiscent of the tree traversal process described by Rosenbaum et al. (1983).

Our work is also intimately related to a broad literature on chunking in sequence learning (Sakai et al., 2003), also referred to as action chunking. Action chunking refers to the “gluing” of consecutive actions that are reinforced repeatedly into a stereotyped action sequence that is executed as a single behavioral unit. One of the most robust findings in the animal learning literature is the emergence of such stereotyped action sequences after extensive training on a particular task. It is thought to occur as control is transferred from a goal-directed system that chooses actions based on their anticipated consequences to a habitual system that executes entire action sequences in response to perceived stimuli (Dolan and Dayan, 2013).

Action chunking has a distinct neural signature, with bursts of neural activity emerging at key choice points as an animal becomes proficient at a particular task (Graybiel, 1998; Barnes et al., 2005; Jin and Costa, 2010). This so-called *task-bracketing* activity first appears in prelimbic cortex – often associated with goal-directed behavior – and then gradually shifts to infralimbic cortex and dorsolateral striatum – often associated with habitual behavior (Smith and Graybiel, 2013). Task-bracketing has also been measured in midbrain dopaminergic neurons that project to striatum (Jin and Costa, 2010), with a difference between the fraction of direct and indirect pathway neurons that code for the initiation and termination of action sequences (Jin et al., 2014). Neural activity representing action sequence boundaries has also been measured in striatum and prefrontal cortex of macaques (Fujii and Graybiel, 2003; Desrochers et al., 2015). Similar task-bracketing activity has also been observed in songbirds (Fujimoto et al., 2011) and humans (Tricomi et al., 2009; Herrojo Ruiz et al., 2014), suggesting a conserved neural mechanism. One interpretation of these results is that task-bracketing activity reflects start/stop signals that gate overtrained action sequences, particularly since it appears to be causally involved in the initiation and termination of action chunks (Geddes et al., 2018).

Executing entire sequences as single behavioral units could be beneficial if the cost of processing the outcome of each action is outweighted by the benefit of acting fast at the risk of making mistakes (Dezfouli and Balleine, 2012). In its current form, our model only captures state chunking (the creation of state clusters), however it can straightforwardly accommodate action chunking using some form of caching or memoization (see Future directions). In fact, our model makes a distinct prediction about the structure of action chunks, namely that they will fall within the boundaries defined by state chunks (Figure 17C). In other words, we predict that state abstraction will drive temporal abstraction: agents will first carve up their environment into clusters of states (state chunks), which in turn will constrain the sequences of actions (action chunks) that are learned, which in turn will operate within the state chunks. This stands in contrast to accounts which assume that the agent first learns useful action sequences and then learns state representations consistent with those sequences post-hoc (Konidaris, 2016). This translates into specific predictions about the neural activity of brain regions thought to support action chunking (see Neural implementation).

### Future directions

Thus far our model only addresses the question of state chunking (how the environment is represented as discrete states at different levels of abstraction), while leaving open the question of action chunking (how actions are stitched together into larger behavioral units at different levels of temporal abstraction). Fitting the model into the HRL framework described above might seem like one way to incorporate action chunking, by assuming that the options leading to bridge endpoints are like action chunks that, when invoked, delegate behavior to a low-level controller that executes the sequence of actions in the option policy as a single behavioral unit. Yet the metaphor of options as action chunks is not necessarily appropriate; option policies execute in a closed-loop fashion, taking into consideration each state they encounter before choosing the next action. In contrast, action chunks are often operationally defined as open-loop action sequences that disregard intermediate states (Dezfouli and Balleine, 2012).

An alternative way to accommodate action chunking that more closely adheres to this definition is to allow caching of solutions to repeated calls to the planner with the same arguments (Huys et al., 2015). That way, when a certain subpath within a cluster is traversed frequently as part of many tasks (for example, getting from your bed your bedroom door), the corresponding action sequences could be cached and, when the subtask needs to be solved as part of a larger task (for example, getting from your bed to work), the action sequence can be retrieved from the cache and executed as a single unit, thus removing the need to unnecessarily recompute it every time. Indeed, such a simple scheme could account for phenomena such as action slips (Dezfouli and Balleine, 2012), or even be used to model task-bracketing activity in striatum (Figure 17C). Implementing action chunking in this way could also account for the speeding up of responses during training in our experiments.

There are several ways in which our model could be extended to support planning in richer, large-scale environments. One way would be to allow deeper hierarchies (*L* > 2), which would be necessary in order to maintain the computational efficiency of the planner as the size of the graph increases. This could be achieved by recursively clustering the high-level states in *H* into higher-level states in another graph *H*′, then clustering those in yet another graph *H*″, and so on up to hierarchy depth *L* that could be prespecified or also inferred from the data. The hierarchical planner can similarly extend recursively by reusing the same logic at the higher levels.

Yet even with deep hierarchies, the learned representations would pose a challenge to the planner as a number of low-level states in *G* grows to the scale encountered in real life (indeed, it would also pose a challenge to the long-term storage system). This highlights another limitation of our model, namely the use of tabular representations in which each state is represented by a separate token. This could be overcome by recognizing that there is redundancy in the environment (Botvinick et al., 2015) – that is, the local structure of the graph *G* can be highly repetitive, with the same theme occurring over and over in different parts of the graph. For example, most cities have streets and buildings, most buildings have rooms and hallways, and most rooms have similar layouts. Representing each and every part of the environment would thus be wasteful, and this redundancy can be exploited by introducing templates or modules – blueprints for clusters that can compress the hierarchical representation by extracting the shared structure across clusters of the same type and only representing differences from some prototypical cluster. Combined with partial observability and deeper hierarchies, this would allow the model to learn representations that support efficient planning in environments that approach the real world in their scale and complexity. Finally, the requirements that *G* is unweighted and undirected can be lifted if the planner is a hierarchical extension of a more sophisticated shortest path algorithm, such as Dijkstra’s algorithm (Cormen et al., 2009).

## Conclusion

In summary, we propose a normatively motivated Bayesian model of hierarchy discovery for planning. The model builds on first principles from the fields of structure learning and robotics, and recapitulates a number of behavioral effects such as detection of boundary transitions, identification of bottleneck states, and surprise to transitions across clusters. The novel predictions of the model were validated in a series of behavioral experiments demonstrating the importance of the task and reward distributions in the environment, which could bias the discovered hierarchy in a way that is either beneficial or detrimental on new tasks. We also showed that the model accounts for reward generalization and uncertainty-based active learning in a way that is consistent with human behavior. Together, these results provide strong support for a computational architecture in which an incremental offline process infers the hidden hierarchical structure of the environment, which is then used by an efficient online planner to flexibly solve novel tasks. We believe our approach is an important step towards understanding how the brain constructs an internal representation of the world for adaptive decision making.

## Acknowledgements

We are grateful to Finale Doshi-Velez, George Konidaris, Bence Ölveczky, Jan Drugowitsch, and Nao Uchida for helpful discussions. This research was supported by the Toyota corporation, the Office of Naval Research (Award N000141712984), the Multi-University Research Initiative Grant (Office of Naval Research Grant N00014-17-1-2961), and the National Institutes of Health (CRCNS 1R01MH109177).

## Appendix

### Inference

We frame hierarchy discovery as computation of the posterior probability distribution *P*(*H*|*D*). Since computing the posterior exactly is intractable, we approximate Bayesian inference over *H* using Metropolis-within-Gibbs sampling (Roberts and Rosenthal, 2009), a form of Markov chain Monte Carlo (MCMC). We initialized *H* by sampling from the prior *P(H*) and on each MCMC iteration, we updated each component of *H* in turn by sampling from its (unnormalized) posterior conditioned on all other components in a single Metropolis-Hastings step, in the following order:

1. update the cluster assignments *c* using the conditional CRP prior (algorithm 5 in Neal, 2000),
2. update *p, q, p′, p″* using a truncated Gaussian random walk with standard deviation 0.1 (i.e. the proposal distribution is a Gaussian centered on the old value, excluding values above 1 or below 0),
3. update update the hierarchical edges *E*′ with a proposal distribution that randomly flips the presence or absence of each edge with probability 0.1,
4. update the average cluster rewards *θ* using a Gaussian random walk with standard deviation 1, and
5. update the average state rewards *μ* using a Gaussian random walk with standard deviation 1.

The samples generated in this way approximate draws from the posterior, with asymptotic convergence to the true posterior in the limit of infinite samples. This approach can also be interpreted as stochastic hill climbing with respect to a utility function defined by the posterior, which has been previously used to find useful hierarchies for robot navigation (Fernández and González, 2013).

In all five simulations and experiments one through seven, for each simulated participant, we generated a Markov chain of a given length (see Table 1) and used the final sample *H* to generate predictions. This can be viewed as a form of probability matching over hierarchies, and is consistent with psychologically plausible algorithms for hypothesis generation and updating, although elucidating the algorithmic details of hierarchy discovery is beyond the scope of our present work.

In order to estimate entropy in experiment eight, we used a separate MCMC sampler for each graph for each participant to generate a set of *M* = 50 samples, using a lag of 100 iterations and burn-in period of 5000 iterations (for a total of 10000 generated samples, as in all other simulations).

Additionally, HBFS requires that the subgraph induced in *G* by each cluster must form a single connected component; that is, for every pair of states (*u, v*) in a cluster *w* = *c_u_* = *c_v_*, there must exist a path (*u*, *x*_1_,…, *x_k_*, *v*) such that *c_x_k__ = w* ∀*k*. To enforce this constraint, we imposed a penalty by subtracting 100 from the (unnormalized) log posterior for each pair of states in the same cluster that are not connected by a path passing through the cluster. This is equivalent to augmenting the generative model with the following rejection sampling procedure: draw *H* according to *P*(*H*|*D*) and for each such pair of disconnected states, perform a Bernoulli coin flip with probability of success equal to *e*^−100^. If all coin flips are successful, keep *H*, otherwise repeat the process with a new *H*. By imposing a “soft” constraint in this way, we ensured that the sampling algorithm can recover from a bad initialization by incrementally adjusting pairs of nodes that violate the constraint. Note that this still results in a valid posterior since normalization is not necessary for approximate inference.

### Decision making

We assume choices based on *H* are either optimal (according to the model) with probably *ε*, or random with probability 1 − *ε*. This is similar in spirit to the *ε*-greedy algorithm in RL, with the difference that we assume the meta-choice to choose randomly occurs before computing the optimal answer. This is equivalent to assuming that on 1 − *ε* of trials, participants simply do not perform the necessary computation. Such lapses could be due to a number of reasons, such as inattention or fatigue. While other factors such as motor variability may also contribute to choice stochasticity, we allow those to be absorbed by the *ε* parameter and leave them as the potential subject of future work.

Another source of variance in choices is the sampled hierarchy *H*, which could be different for each simulated participant. While our current experiments prohibit direct comparisons between hierarchies inferred by participants and hierarchies inferred by the model, any systematic variation in the sample hierarchies (for example, due to uncertainty of the posterior, as in experiments three and eight) would manifest when choices are aggregated across participants, as in our analyses.

#### Hierarchical planning

The optimal algorithm to find the shortest path between a pair of low-level vertices (*s, g*) in *G* is breadth-first search (BFS; Cormen et al., 2009) whose time and memory complexity is *O*(*N*) (assuming *O*(1) vertex degrees, i.e. |*E*| = *O*(*N*)). We use a natural extension of BFS to hierarchical graphs (hierarchical BFS or HBFS; Fernández and González, 2013) that leverages *H* to find paths more efficiently than BFS (approximately 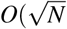 time and memory). Intuitively, HBFS works by first finding a high-level path between the clusters of *s* and *g*, *c_s_* and *c_g_*, and then finding a low-level path within the cluster of *s* between *s* and the first bridge on the high-level path.

In particular, HBFS first finds a high-level path (*w*_1_,…, *w_m′_*) between *c_s_* and *c_g_* in the high-level graph *H* (note that *w*_1_ = *c_s_* and *w_m′_* = *c_g_*). Then it finds a low-level path (*y*_1_,…, *y_m_*) between *s* and *u* in *G*[*S*] (note that *s* = *y*_1_ and *u* = *y_m_*), where (*u, v*) = *b*_*w*_1__, *w*_2_ is the first bridge on the high-level path, *S* = {*x*: *c_x_* = *c_s_*} is the set of all low-level vertices in the same cluster as *s*, and *G*[*S*] is the subgraph induced in *G* by *S*. HBFS then returns *y*_2_, the next vertex to move to from *s*, or, alternatively, full the path to the next cluster, (*y*_2_,…, *y_m_*).

In an efficient hierarchy, the number of clusters will be 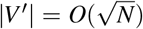 and the size of each cluster *w* will also be 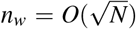, resulting in 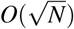 time and memory complexity for HBFS. Note that actually traversing the full low-level path from *s* to *g* in *G* still takes *O*(*N*) time; HBFS simply computes the next step, ensuring the agent can progress towards the goal without computing the full low-level path in advance (in our simulations, we actually computed the full path in order to simulate execution in addition to planning). HBFS can straightforwardly extend to deeper hierarchies with *L* > 2, with the corresponding complexity becoming 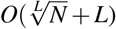.

##### Algorithm 1 HBFS(*s, g, H, G*)

1: *path′* ← BFS(*c_s_*, *c_g_*, (*V*′, *E*′))

2: *path* ← []

3: **for all** (*w,z*) in *path*’ **do**

4: (*u, v*) ← *b_w,z_*

5: *S* ← {*x*: *c_x_* = *c_s_*}

6: append (*path*, BFS(*s, u, G*[*S*]))

7: append(*path*, (*u, v*))

8: *s* ← *v*

9: **end for**

10: *S* ← {*x*: *c_x_* = *c_s_*}

11: append(*path*, BFS(*s,g, G*[*S*]))

12: **return** *path*

The pseudocode for HBFS used in our simulations is shown in Algorithm 1. Our particular implementation of HBFS takes as arguments the starting state *s* ∈ *V*, the goal state *g* ∈ *V*, the hierarchy *H* and the low-level graph *G*. The variables *c* and *b* refer to the cluster assignments and bridges in *H*, respectively. Note that *s* changes after each iteration. We assume the existence of a function BFS which takes as arguments a starting state, a goal state and a graph and returns the shortest path between those states as a list of edges.

Note that we are not making any specific commitments to the cognitive plausibility of HBFS, and that any other hierarchical planner based on shortest paths would make similar predictions. Also note that HBFS could be straightforwardly extended to deeper hierarchies by introducing a depth level parameter *l* and recursively calling HBFS instead of BFS on line 1 if *l* > 2. Finally, note that HBFS as implemented here still requires *O*(*N*) time and memory as it finds the full path in *G*. For returning only the first few actions, the for-loop could be interrupted after the first iteration, which would yield the 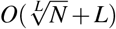 complexity.

#### Rewards

To model reward generalization in experiment six, we set 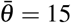 and *r*_4,1_ = 30, in accordance with the experimental instructions. The node selected by the simulation was the one with the greater (approximate) expected reward 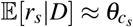, where *θ* are the cluster rewards and *c* are the cluster assignments in the sampled hierarchy *H*.

To model cluster inferences based on rewards in experiment seven, we only simulated a single training trial and set *r*_1,1_ = *r*_2,1_ = *r*_3,1_, *r*_4,1_ = *r*_5,1_ = *r*_6,1_, and *r*_7,1_ = *r*_8,1_ = *r*_9,1_ = *r*_10,1_, with the specific values chosen at random between 0 and 30 (we scaled down the rewards experienced by participants by a factor of 10). Since the hierarchy discovery algorithm has no notion of maximizing reward (it merely treats rewards as features), the magnitude of the rewards is irrelevant. We did not model the random changes of reward values throughout training, which were introduced purely for control purposes and are irrelevant for hierarchy discovery as framed here. Note that the model could easily accommodate dynamic rewards by assuming drifting *μ*’s, however this would unnecessarily complicate the model without making substantial contributions to the core theoretical predictions. As in experiment six, we set 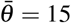, the expected reward based on the instructions.

#### Active learning

Drawing on the active learning framework from the causal inference literature (Murphy, 2001; Tong and Koller, 2001), we assume the agent will chose to learn about edges of *G* in a way that provides maximal information about *H*. Maximizing information about *H* is equivalent to minimizing uncertainty about *H*, which can be quantified as the entropy of *H*:

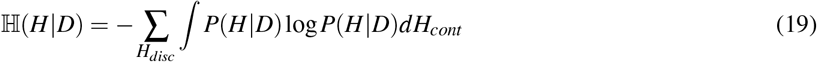

Where *H_disc_* = (*V*′, *E*′, *c*, *b*) are the discrete components of *H*, *H_cont_* = (*p*′, *p*, *q*) are the continuous components of *H*, and *D* is the data observed so far. We use 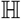 to denote the entropy of a mixed random variable with discrete and continuous components (Nair et al., 2006).

Computing the entropy in this way is neither computationally feasible nor psychologically plausible. Following previous authors (Steyvers et al., 2003), we assume the agent has a subjective probability distribution over possible hierarchies *H* which can be represented by a set of samples [*H*^(1)^,…, *H*^(*M*)^] with multinomial probabilities 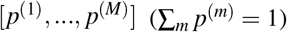. If the samples are drawn from the posterior *P*(*H*|*D*), the agent can approximate the entropy as:

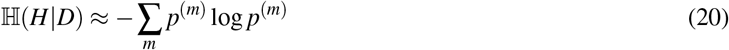

Note that while this is not a proper estimate of the entropy, it can serve as a basis for rational hypothesis testing. In our simulations, we used MCMC to generate the samples from the (approximate) posterior and set the subjective probabilities according to *p*^(*m*)^ ∝ *P*(*H*^(*m*)^|*D*) ∝ *P*(*D*|*H*^(*m*)^)*P*(*H*^(*m*)^).

We use *a_u,v_* to denote the action of observing edge (*u, v*), i.e. finding out whether (*u, v*) ∈ *E*. Since there is no way to know in advance what the outcome would be, the agent has to minimize the expected entropy over the two possible outcomes:

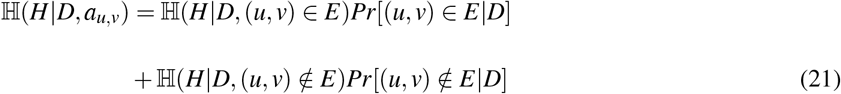

We can compute the probability of each outcome by marginalizing over *H* and using the sampling approximation:

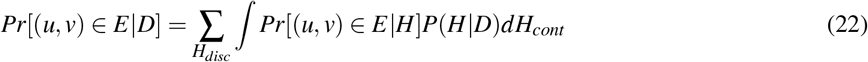

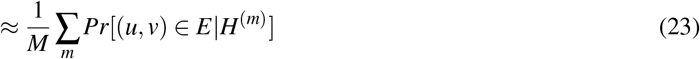

Where *Pr*[(*u, v*) ∈ *E*|*H*] is *p* or *pq*, according to the generative model. *Pr*[(*u, v*) ∉ *E*|*D*] is approximated analogously.

The agent then chooses the action that minimizes the expected entropy:

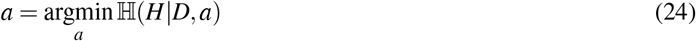

### Neural simulations

The example within-trial circuit activations in Figure 17B were generated manually, assuming a hierarchy like the one in Figure 17A (respondng to the decomposition in Figure 9A). We assumed HBFS is executed at every cluster boundary and that the entire path within the cluster is traversed in a single action sequence that is executed as a single behavioral unit, akin to an action chunk.

To generate the Bayesian update in Figure 17D, we simulated online inference on the graph from the Towers of Hanoi puzzle (Figure 7A), using a particle filter with *M* = 100 particles [*H*^(1)^,…, *H*^(*M*)^], each initialized from the prior *P*(*H*). We started with an empty graph and added edges one by one, with single edge added on each trial. We approximated the posterior *P*(*H*|*D*) with multinomial probabilities [*p*^(1)^,…, *p*^(*M*′)^], where *p*^(*m*)^ ∝ *P*(*H* ∝ *P*(*D*|*H*^(*m*)^)*P*(*H*^(*m*)^) and 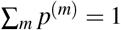.

Following our previous work (Tomov et al., 2018), we quantified the Bayesian update after each observed edge by computing the Kullback-Liebler divergence between the multinomial approximation to the posterior before and after the update. We additionally performed 10 iterations of MCMC (as described in the inference section) for each particle after each trial in order to rejuvenate the particles (Chopin, 2002). Similar approximations to online Bayesian inference have been used in previous studies (Abbott and Griffiths, 2011). To generate the dissimilarity matrix in Figure 17E, as in our previous work (Tomov et al., 2018), we computed the cosine distance between the approximate posterior for each pair of trials in the simulation.

Data and code for all simulations and experiments are freely available at https://github.com/tomov/chunking.

